# CAMDA 2023: finding patterns in urban microbiomes

**DOI:** 10.1101/2024.09.26.614772

**Authors:** Haydeé Contreras-Peruyero, Imanol Nuñez-Morales, Mirna Vazquez-Rosas-Landa, Daniel Santana-Quinteros, Antón Pashkov, Mario E. Carranza-Barragán, Rafael Perez-Estrada, Shaday Guerrero-Flores, Eugenio Balanzario, Víctor Muñiz Sánchez, Miguel Nakamura, L. Leticia Ramírez-Ramírez, Nelly Sélem-Mojica

## Abstract

The Critical Assessment of Massive Data Analysis (CAMDA) addresses the challenge of effectively utilizing Big Data in life science. Serving as both a conference and a catalyst for research groups, CAMDA annually presents challenges that foster innovative solutions. For the Forensics CAMDA 2023 challenge, we analyzed 365 metagenomic samples from 16 cities worldwide to characterize their origin. The forensic challenge was addressed from two perspectives: using the reduced abundance OTU tables and employing functional annotations. To identify the most informative Operational Taxonomic Units, we fit negative binomial models ultimately reducing variables to 294. After OTU selection, we implemented supervised models and conducted 5-fold cross-validation (CV) with a 4:1 training-to-validation ratio in each scenario. Support vector classification (SVC) achieved the highest F1 score (0.96) for the abundance tables, accurately classifying most cities, although New York City (NYC) posed a challenge. Via functional profiles with Mifaser at level 4, we achieved the best functional classification using the Neural Network (NN) model.

Additionally, to gain insight into further associations between bacterial distribution with other covariates, we applied Dirichlet regression over *Escherichia, Enterobacter*, and *Klebsiella* bacteria abundances. We considered climatic and demographic variables of the cities, observing that population increase is indeed associated with a rise in the mean of *Escherichia* while decreasing temperature is linked to higher proportions of *Klebsiella*.

For replicability of the scripts, a Docker container and a Conda environment are available at the repository: GitHub:github.com/ccm-bioinfo/cambda2023.

## 1 INTRODUCTION

The Critical Assessment of Massive Data Analysis (CAMDA) addresses a fundamental challenge of our era: the effective and intelligent utilization of Big Data in the life sciences. Microbes constitute most of Earth’s biodiversity (Hug et al., 2016); however, our comprehension of their distribution and ecological roles remains incomplete. Urban microbiomes, in particular, offer valuable insights into living conditions and public health. Studying the microbiome of public transportation systems can be a proxy for monitoring other urban sites. Various factors influence the presence and metabolic content of microorganisms in urban environments. The microbiome of public transportation systems is largely derived from the human microbiome (Hernández et al., 2020). A primary source of variation in the human microbiome is the specific niche (Huttenhower et al., 2012). The microbiome found in public transportation systems is reflective of the skin microbiome. Although the skin microbiome can vary over time (Gilbert et al., 2018), some studies indicate a degree of stability (Byrd et al., 2018; Callewaert et al., 2020). After traveling, passengers’ microbiomes tend to converge (Vargas-Robles et al., 2020). Subways act as hubs for microbiome exchange and are reservoirs for antibiotic resistance (Peimbert and Alcaraz, 2023). The forensic challenge aimed to develop models capable of accurately classifying these metagenomic samples according to their city of origin.

Since 2017, the CAMDA community has promoted forensic metagenomics challenges in collaboration with the MetaSUB consortium. In the 2017 challenge, approximately 1000 samples of 16S from three cities in the United States suggested that differences in the cities’ microbiome were enough to separate them (Walker et al., 2018). For the CAMDA 2018 challenge, 293 shotgun metagenomics samples from 12 cities were analyzed, extending the 16S data from 2017 (Walker and Datta, 2019). In 2018, benchmarking of genome ensemble methods from MetaSUB metagenomes was conducted (Gerner et al., 2018). In 2019, participants were interested in functional annotation, including antibiotic resistance (Casimiro-Soriguer et al., 2019), and even a microbiome annotator was developed (Zhu et al., 2017, 2019). In 2020 challenge, 28 cities were analyzed (Zhang et al., 2021a,b). Other variables such as climate and population density were included (Zhelyazkova et al., 2021) and model combinations were included (Walker et al., 2018; Anyaso-Samuel et al., 2021b,a). Finally, in 2021, outside of CAMDA challenges, a global map of the urban microbiome and antibiotic resistance was created by Danko, complete with a web interface (Danko et al., 2021) featuring more than 4000 samples from 60 cities. In 2023, CAMDA presented a classification challenge involving microbiomes from public transportation systems in 16 cities worldwide. A total of 365 metagenomic samples were provided to the CAMDA community by The International Metagenomics and Metadesign of Subways and Urban Biomes (MetaSUB) Consortium (Mason et al., 2016), see table 1 1 for a complete description of the samples and variables provided (city, year, and number of samples) and collected (latitude, longitude, climate, average of reads, and standard deviation)”.

**Table 1.**
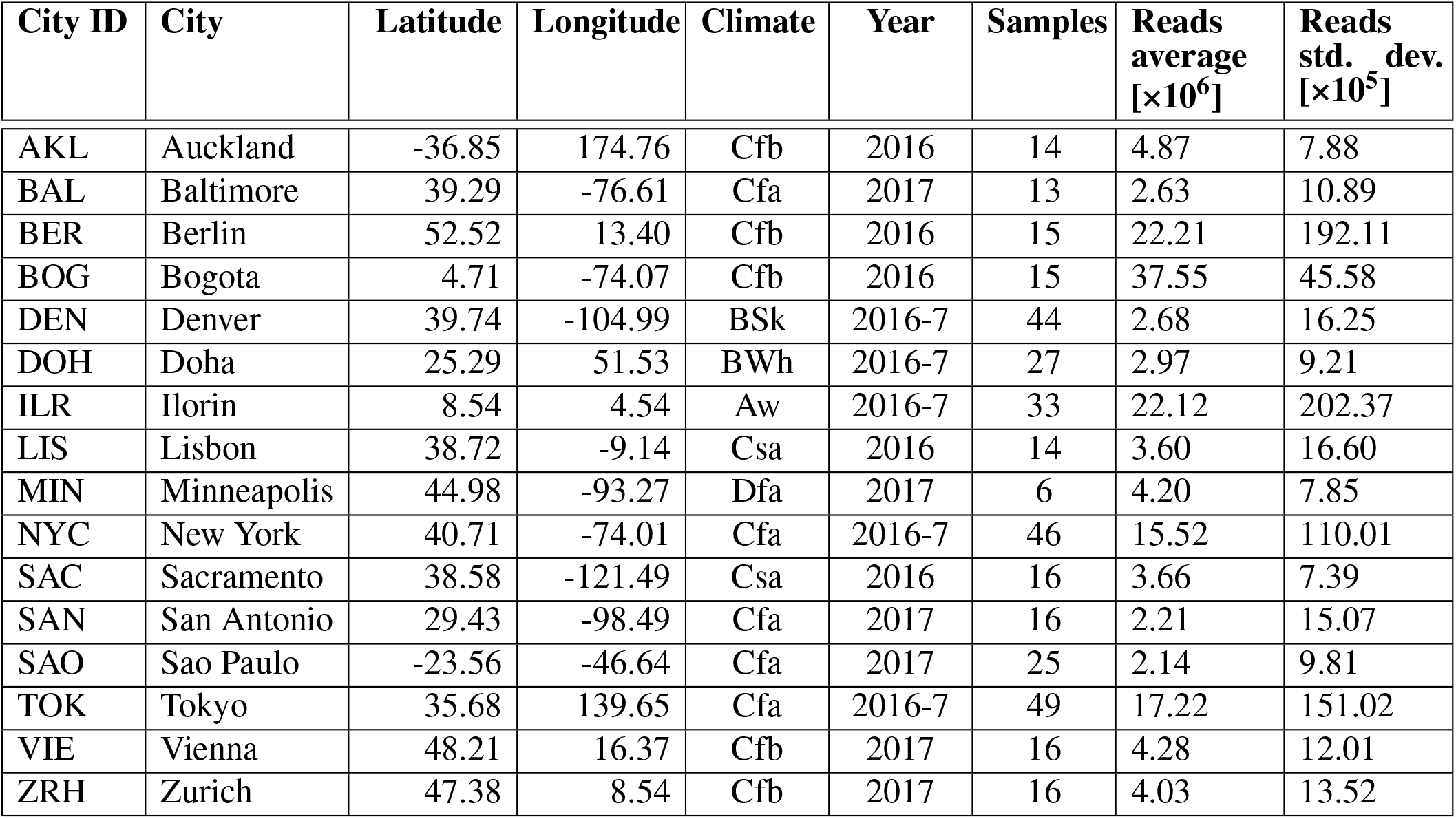
Summary table of the cities from which metagenomic samples were collected. The city set covers multiple places around the world and different types of climates. Climate symbols follow Köppen climate classification. Samples were collected in 2016, 2017, or both (marked as 2016-7 in the table). The cities were not sampled equally: there are significant differences in the sample count and read depths between cities. For example, Tokyo and New York lead in terms of sample count, whereas Minneapolis has the least samples. Bogota was sampled with a significantly higher depth than other cities, while samples from San Antonio and Sao Paulo have a relatively low read depth.

| City ID | City | Latitude | Longitude | Climate | Year | Samples | Reads average<br>[$\times 10^6$] | Reads std. dev.<br>[$\times 10^5$] |
| --- | --- | --- | --- | --- | --- | --- | --- | --- |
| AKL | Auckland | -36.85 | 174.76 | Cfb | 2016 | 14 | 4.87 | 7.88 |
| BAL | Baltimore | 39.29 | -76.61 | Cfa | 2017 | 13 | 2.63 | 10.89 |
| BER | Berlin | 52.52 | 13.40 | Cfb | 2016 | 15 | 22.21 | 192.11 |
| BOG | Bogota | 4.71 | -74.07 | Cfb | 2016 | 15 | 37.55 | 45.58 |
| DEN | Denver | 39.74 | -104.99 | BSk | 2016-7 | 44 | 2.68 | 16.25 |
| DOH | Doha | 25.29 | 51.53 | BWh | 2016-7 | 27 | 2.97 | 9.21 |
| ILR | Ilorin | 8.54 | 4.54 | Aw | 2016-7 | 33 | 22.12 | 202.37 |
| LIS | Lisbon | 38.72 | -9.14 | Csa | 2016 | 14 | 3.60 | 16.60 |
| MIN | Minneapolis | 44.98 | -93.27 | Dfa | 2017 | 6 | 4.20 | 7.85 |
| NYC | New York | 40.71 | -74.01 | Cfa | 2016-7 | 46 | 15.52 | 110.01 |
| SAC | Sacramento | 38.58 | -121.49 | Csa | 2016 | 16 | 3.66 | 7.39 |
| SAN | San Antonio | 29.43 | -98.49 | Cfa | 2017 | 16 | 2.21 | 15.07 |
| SAO | Sao Paulo | -23.56 | -46.64 | Cfa | 2017 | 25 | 2.14 | 9.81 |
| TOK | Tokyo | 35.68 | 139.65 | Cfa | 2016-7 | 49 | 17.22 | 151.02 |
| VIE | Vienna | 48.21 | 16.37 | Cfb | 2017 | 16 | 4.28 | 12.01 |
| ZRH | Zurich | 47.38 | 8.54 | Cfb | 2017 | 16 | 4.03 | 13.52 |

Reducing the predictor variables to a small set of OTUs (Ryan, 2019; Casimiro-Soriguer et al., 2019; Zhang et al., 2021a), functions, or AMR features that constitute a footprint of a city has been a constant goal in the CAMDA challenges. The Negative Binomial (NB) model is a generalized linear model suited for counting overdispersed data. Since the work of Lu et al. (Lu et al., 2005), the NB model has found extensive application in the analysis of differential gene expression, see for instance the R packages edgeR (Chen and Lun, 2017) and DeSeq2 (Michael Love, 2017). Drawing inspiration from this application, McMurdie et al. (McMurdie and Holmes, 2014) introduced the NB model for microbiome count data analysis to identify differentially abundant OTUs. In this work, we apply the NB model to guide variable selection as a first step before classification algorithms. Specifically, we employ NB to identify differentially abundant OTUs across multiple cities, aiming to reduce data dimensionality. Following data dimension reduction, we proceeded to evaluate the performance of several classification algorithms on our dataset. The best result was obtained with Support Vector Classifier (SVC), which achieved an F1 score of 0.96, followed by Multilayer Perceptron Classification (MLPC) and K-Nearest Neighbors (KNN), yielding scores of 0.95 and 0.91, respectively.

In addition to the challenges of classification, the community has been interested in understanding the relationship between microbiomes and other variables, such as climate, population density of the city, or the specific surface of the transport where the sample was collected. This year, CAMDA highlights three clinically significant bacteria: *Klebsiella, Escherichia*, and *Enterobacter*, in addition to microbiome data, which consisted of urban microbiome samples from 16 cities in the world, these samples are collected every year on June 21. Dirichlet regression (Maier, 2014) is employed to understand the statistical distribution of relative abundances depending on other variables known as covariates. This study focuses on these three bacteria of interest and examines how their abundances relate to a set of climatic and demographic variables associated with the samples’ dates of acquisition. In contrast to Zhang et al.’s (Zhang et al., 2021b) emphasis on city-specific climate metadata, our approach integrates demographic factors and finds them relevant. Danko et al. (Danko et al., 2021), consider climate-related, demographic and geographical components and discard population density in their context, whereas we find that population density emerges as a statistically significant factor related to microbial composition.

Forensic metagenomics has demonstrated the ability to differentiate the origin city of a microbiome from the collective transport system for several years (Walker et al., 2018; Walker and Datta, 2019; Danko et al., 2021). This study used the NB model to select taxonomic variables before implementing supervised algorithms to differentiate microbiomes by city. Then we incorporate functional profiles, with various annotation methods, where Mifaser at level 4 provided the best performance with MLP, achieving an F1 score of 65.4%, followed by Metacyc at level 7 with VC(Soft) and an F1 score of 52%. Finally, we **conducted a Dirichlet analysis and found that there is an impact of other variables, such as climate and** population density, on the abundance of these bacteria.

## 2 RESULTS

The objective of the forensic geolocalization challenge is to predict the city of origin for selected samples using taxonomic and functional profiles. The MetaSUB consortium provided CAMDA participants with urban microbiome samples from 16 cities worldwide. Each year, on June 21, the Global City Sampling Day (gCSD) collects microorganisms in urban environments. From the numerous samples collected during gCSD 2016 and 2017, 374 paired-end samples were made available for this challenge (Figure 1a). Following quality control, nine samples were excluded from the analysis. The remaining 365 samples—174 from 2016 and 191 from 2017—underwent adapter and quality trimming, reducing their total size by approximately 30%, and were subsequently assembled. Using both the read and contigs, we constructed their taxonomic profiles, identifying around 20,000 OTUs among all reads and nearly 13,000 in the assemblies distributed across the 16 cities (refer to Figure 1b for the complete pipeline). Apart from Homo sapiens, with a read proportion of 33.35%, the most abundant taxa were *Cutibacterium acnes* (3.93%), *Stutzerimonas stutzeri* (3.85%), *Bradyrhizobium sp. BTAi1* (1.96%), *Massilia sp. NP310* (0.80%) and *Staphylococcus aureus* (0.67%), all of which correspond to human or soil-associated bacteria; see Figure S1.

**Figure 1.**
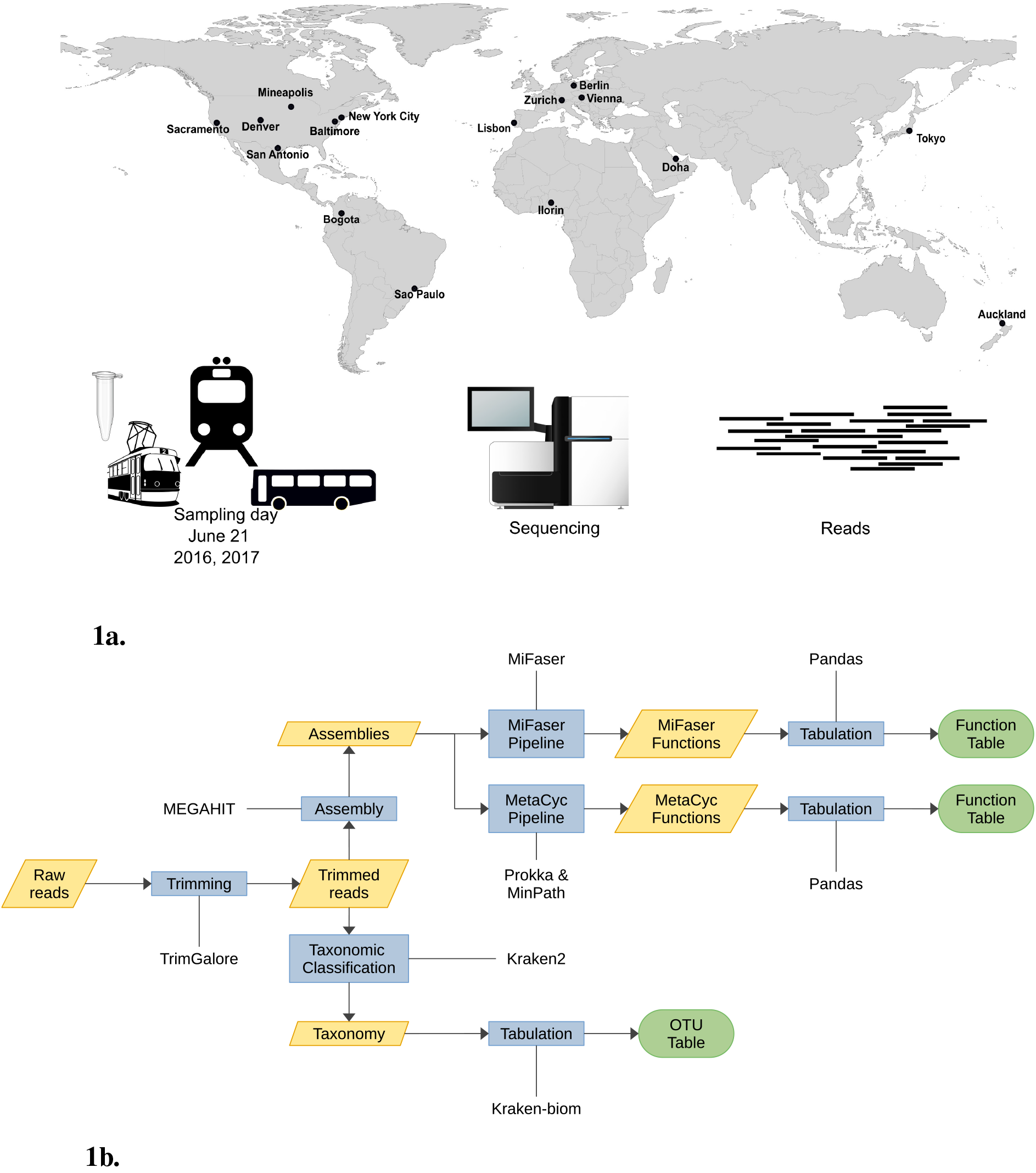
**(a)** On June 21th every year MetSUB collected urban samples around the world. For the CAMDA 2023 challenge, we were provided 374 samples from 16 cities. **(b)** The 374 samples underwent prepocessing. Nine samples were discarded in the initial quality control step, leaving 365 samples for the adapter and quality trimming process. During this phase, their size was reduced by 30% before assembly. Subsequently, both reads and assemblies were used for taxonomic assignment.

As an initial city classification model, we utilized principal components followed by a multivariable logistic regression. First, we selected a subset of *J* OTUs from a total pool of 20,448. After applying a natural logarithm transformation, this subset underwent principal component analysis (PCA), generating the first *N* principal components to serve as inputs for the logistic regression classification algorithm. Various combinations of *J* and *N* were tested, achieving accuracy of up to 88% in the classification task. Since increasing the value of *J* did not significantly improve performance, we applied variable selection on the reads table, followed by several classification models, refer to the Supplementary Material Section 3 for a full description of the method. As a result, fewer than 300 variables were chosen based on their status as the most differential OTUs under a Negative Binomial model.

### 2.1 Selecting most differential OTUs

We aimed to strategically curate Operational Taxonomic Units (OTUs) that best differentiate by city and year, considering potential zero-inflated OTU distribution. By modeling the zero-inflated data phenomenon in our selection process, we aimed to impact the reliability and effectiveness of our predictive frameworks. The proposed models were fitted for the data organized by reads and assembly, each categorized by Bacteria-Archaea, Eukarya, Virus, and all three categories combined. We further divided each of the eight resulting datasets, see Table 2, to consider their components at different taxonomic levels: phylum, class, order, family, and genus, resulting in 40 databases. This comprehensive approach ensures a thorough examination of the data.

**Table 2.** Datasets obtained by considering reads and assembly data, categorized by Bacteria-Archaea, Eukarya, and Virus. The column Dataset describes the name of each dataset, while the ✓ mark indicates what data is contained in it.

| Dataset | Type of data |  | Categories |  |  |
| --- | --- | --- | --- | --- | --- |
|  | Reads | Assembly | Bacteria-Archaea | Eukarya | Virus |
| reads | ✓ | — | ✓ | ✓ | ✓ |
| readsAB | ✓ | — | ✓ | — | — |
| readsEukarya | ✓ | — | — | ✓ | — |
| readsViruses | ✓ | — | — | — | ✓ |
| assembly | — | ✓ | ✓ | ✓ | ✓ |
| assemblyAB | — | ✓ | ✓ | — | — |
| assemblyEukarya | — | ✓ | — | ✓ | — |
| assemblyViruses | — | ✓ | — | — | ✓ |

Since selecting important variables for predictive models is reasonably related to identifying differentially abundant OTUs, similarly to the DESeq2 library from R (Love et al., 2014), we consider generalized linear models for count data 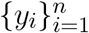, but including models that can account for a possible excess of zeros in the data. The four models were Poisson, Negative binomial (NB), Zero-inflated Poisson (ZIP), and Zero-inflated negative binomial (ZINB). In all of them, covariables {*x*_*i*_} and {*z*_*i*_} represent a pair city-year, see Methods 4.4. Specifically, given two city-years, {*x*_*i*_} and {*z*_*i*_} are dummy variables that indicate the city-year to which the counts {*y*_*i*_} are associated. For each OTU and each categorical pair of variables {*x*_*i*_} and {*z*_*i*_}, *p*-values were collected for the coefficients of the fitted regressions to assess their statistical significance with False Discovery Rate adjustments. Afterward, within each model, we selected OTUs with the lowest associated adjusted *p*-values for each combination of year and city.

As we simultaneously conduct multiple hypothesis tests on various Operational Taxonomic Units (OTUs), the probability of committing a Type I error increases with the number of tests. This error is related to the ‘False Discovery Rate’ (FDR), and it occurs for a specific pair of variables (year-cities in our case) when a small *p*-value leads to the rejection of *H*_0_: ‘There is no differential OTU count between the compared year-cities,’ when *H*_0_ is actually true (that is, there is no statistical difference between the OTU counts of the two year-cities). The FDR is the expected proportion of false positive results among all the rejected hypotheses. Various methods can be employed to mitigate this tendency and prevent spurious discoveries. One of the most well-known methods is the Bonferroni correction (Bonferroni, 1935), where the base significance level is divided by the number of tests. Alternatively, more sophisticated approaches, such as the Tukey (Tukey, 1949) or Scheffé tests (Scheffé, 1999), can be utilized. Here, to mitigate incorrect rejections of the true null hypothesis, i.e., to control the FDR, we used the Benjamini-Hochberg procedure (Benjamini and Hochberg, 1995).

We observed that many *p*-values were exceptionally low (numerically equal to zero) but presented high Akaike information criterion (AIC) scores, pointing to low model fitting. The AIC is a measure used in statistics and model analysis to compare how suitable each model is; the lower its value, the better the model fits. Thus, for each OTU and each pair of year-cities, we selected the model with the minor AIC score and kept the *p*-value associated with that model. This prevents the presence of OTUs associated with small *p*-values but with inappropriate fitting. After computing the AIC scores, we concluded that NB model was the most appropriate fitting. Then we proceed to choose differential OTUs. We deemed an OTU to be “most differential” if it is differential for more than ten year-city comparisons, i.e., if the OTU is selected as a differential for at least 11 pairs of year-cities. The table of the most differential OTUs is presented in the Supplementary material, Table S1.

The NB model, fitted on each kingdom separately, was more successful in selecting the 294 most informative OTUs (see Figure 2a for the complete pipeline, and Table 3 for the number of selected OTUs when considering the 5 OTUs with lowest *p*-values). Considering the NB model and individual kingdoms to select variables (differential OTUs), classification reached an F1 score close to 0.96, see the results described in section 2.2. Pairwise comparison and selection of most differential OTUs might be interpreted biologically. Alpha diversity measures environmental richness or species abundance within an environment. In Figure 2b, we can observe the richness distribution in each sample. After implementing the selection process for the most informative Operational Taxonomic Units (OTUs), we observe that the distribution of alpha diversity remains consistent across each sample, Figure 2c.

**Table 3.** Number of selected OTUs under each model. In parenthesis we indicate the percentage of OTUs that were not selected in any pair of year-cities.

| Model | Fit with all kingdoms | Fit kingdoms separately |
| --- | --- | --- |
| P | 15 (0.91) | 16 (0.96) |
| NB | 280 (8.19) | 294 (10.24) |
| ZIP | 343 (11.85) | 305 (10.70) |
| ZINB | 450 (29.94) | 428 (30.22) |
| Model selection | 298 (12.76) | 312 (11.94) |

**Figure 2.**
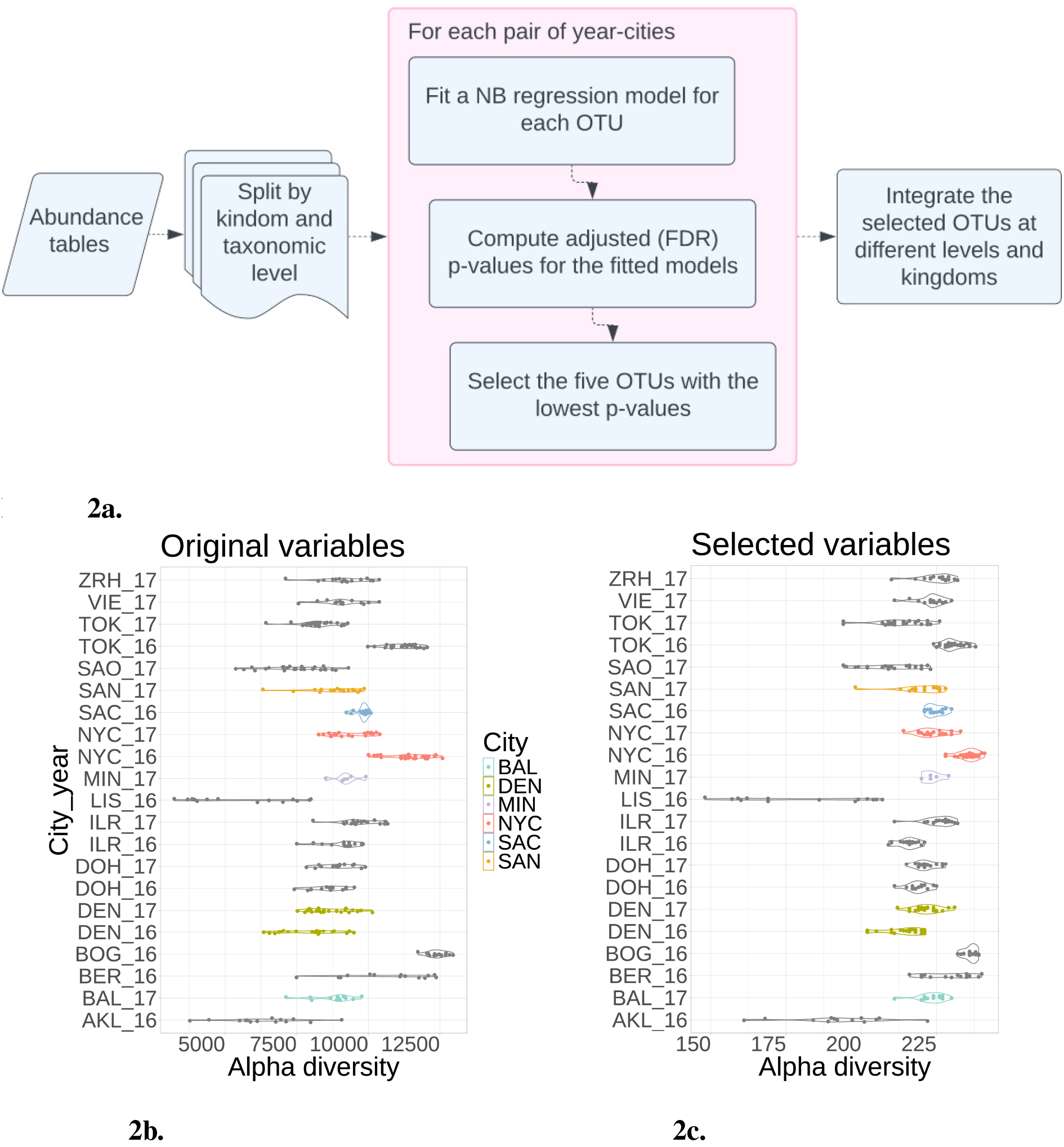
Negative Binomial model reduced variables by selecting 300 from 20,000 OTUs. **(a)**Pipeline to fit the NB model in the abundance tables. **(b)** Alpha diversity from each city and year considering all the OTUs. **(c)** After selecting the most distinguishable OTUs, the alpha diversity maintains a consistent distribution across each city. This suggests that reducing variables preserves the original samples’ richness.

### 2.2 Prediction Models using OTUs selection

Classification models were implemented with a 5-fold cross-validation scheme with 4:1 training to validation sets. OTUs were selected using the NB selection method, independently for each of the 5 folds in the cross-validation scheme as described in Figure 3c. This ensured that no information from one fold was used to influence the classification for the test set, thereby avoiding model overfitting or model score bias. In this process each fold got between 288 and 304 different OTUs with 123 of them being shared between all folds, detailed description of features (OTUs) used for each fold presented in Supplementary Material table 1.2. Macro F1 scores for city prediction models based on abundance tables were computed. The Multilayer Perceptron Classifier (MLPC) achieved a score of 0.95, the Support Vector Classifier (SVC) scored 0.96, and the K-Nearest Neighbor (KNN) model attained a score of 0.91 (rounded values). The results presented in Figure 3a illustrate the consistency of these scores across different folds, with the SVC model exhibiting less variance in the Macro F1-score compared to KNN and MLPC, which reported broader confidence intervals. Furthermore, Figure 3b, shows the outstanding accuracy of the SVC model in predicting most cities.

**Figure 3.**
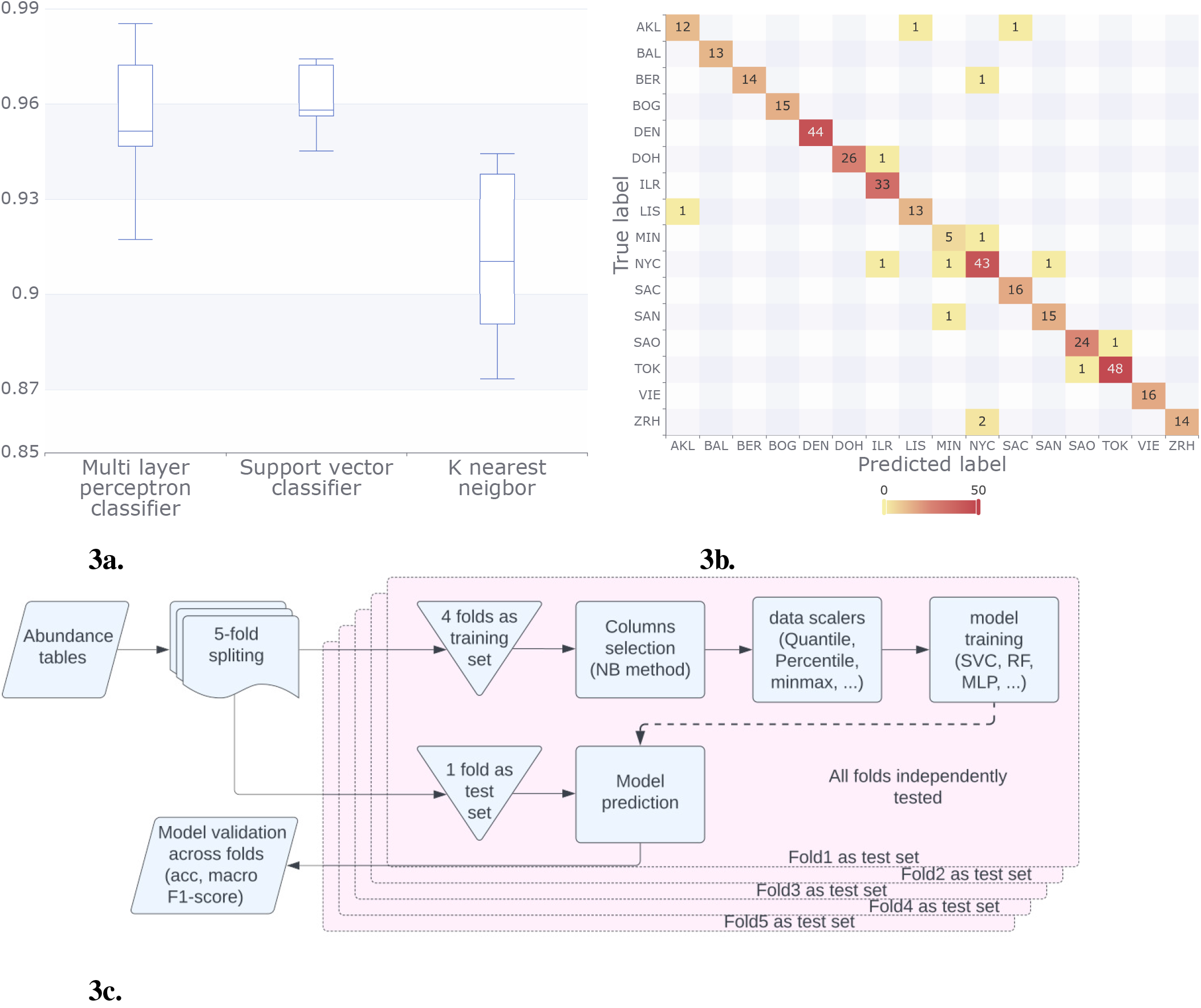
Results for the classification methods considering the abundance tables. **(a)** For each classification model, we plotted the Macro F1 score, in all the model this score was above 0.9. **(b)** In the confusion matrix for the SVC model, we can observe that the majority of predictions were correct (numbers on the diagonal). NYC had the most incorrect labels (numbers off the diagonal). **(c)** We split the data (abundance tables) into five folds; after this, we trained the models using four-folds, and we tested our result with the remaining fold. We run this process by varying the folds in the training set and the validation set. Finally, we compared the acc and made F1 scores of the models.

### 2.3 Classification models with functional profiles

We annotated the functional profiles within our samples with the tools: Mifaser(Zhu et al., 2017), Metacyc (Caspi et al., 2019) and Prokka(Seemann, 2014a). In the case of Mifaser and Metacyc, we kept data structured into hierarchical tables based on the level of functional annotation. For Mifaser, annotations are E.C. numbers stored in data tables ranging from level 1 to level 4, where level 4 provided the highest annotation specificity. Similarly, with Metacyc, data tables span specificity levels from 1 to 8, with level 8 offering the most detailed annotations.

For instance, a specificity level 1 annotation in Mifaser could be as broad as “oxidoreductases (EC 1)” encompassing a wide range of enzymes that catalyze oxidation-reduction reactions. A more specific level 4 annotation in Mifaser might include terms like “glycerol-3-phosphate dehydrogenase (EC 1.1.1.8)” which provides a more detailed description of the enzyme’s function. On the other hand, in MetaCyc, a specificity level 1 function might be broadly categorized as “biosynthesis” and a more specific level 8 annotation could encompass detailed pathways such as “CMP-legionaminate biosynthesis I”.

We used a similar scheme to that employed with abundance tables, implementing a 5-fold cross-validation scheme with 4:1 training to validation sets. However, unlike the approach with abundance tables involving the NB model, we utilized a quantile transformer before training the models, Figure 4c.

**Figure 4.**
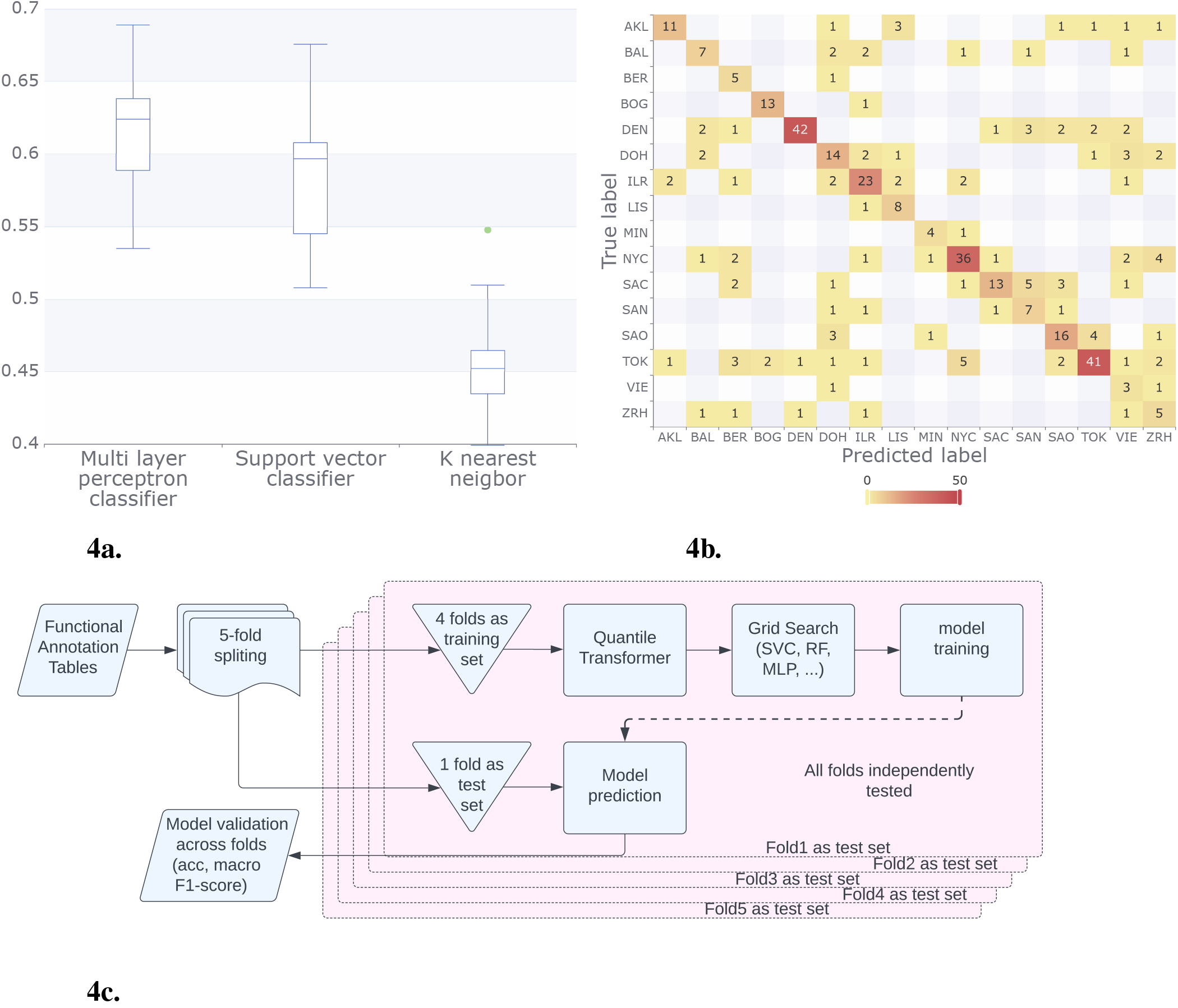
Results for the classification methods considering the functional annotation. **(a)** Using Mifaser functional annotation tables level 4, the trained models showed improved and consistent performance, achieving a macro F1 score between 0.4 and 0.6. The MLP model exhibited the best performance on average. **(b)** The confusion matrix for the MLP model shows that some cities reached up to 92% correct responses, with BAL, SAC, and ZRH being the lowest. **(c)** In contrast to the pipeline that utilizes abundance tables, for the functional annotation tables we employed a quantile transformer after implementing a 4:1 training to validation set scheme. This approach allowed us to sweep parameters and train the models effectively.

Our analysis assessed the performance of the following machine learning models: KNN, MLP, and SVC. Additionally, we explored alternative methods and their ensembles. The supplementary materials provide detailed results and findings from these additional analyses. It is important to note that the results in the supplementary materials did not include a 5-fold cross-validation.

The model results consistently demonstrated superior performance when utilizing Mifaser data at specificity level 4 where, after a rigorous 5-fold cross-validation process, we found that the Multi-Layer Perceptron (MLP) Classifier demonstrated superior performance. With a mean macro F1 of approximately 60% across the folds as shown in Figure 4a

### 2.4 Dirichlet regression reveals associations with concomitant variables

Regression methods characterize the statistical distribution of a relevant variable (called the response) as a function of other variables that exert some influence or association (called predictors or covariates). The expectation is that discovering the relationships between these sets of variables sheds fascinating biological insight. We conceive regression in its broadest sense: describe the complete distribution of the response instead of only its mean. This year, beyond the microbiome data, CAMDA focuses on three clinically relevant bacteria. As the response, we consider the relative abundance of our three bacteria of interest, *Klebsiella, Escherichia*, and *Enterobacter*. This constitutes a case of *compositional data*, meaning that each observed city sample datum is a set of three proportions that add up to 100%. The Dirichlet distribution is a flexible statistical model specifically suitable for this type of observation.

Studies on climate change have shown that temperature and humidity variations impact the composition, structure, regulation, and presence of metabolic functions in microbial communities (Jansson and Hofmockel, 2020). The climatic effect depends on the ecosystem, and population density can also be significant in urban environments. For each city, we augmented its original genomics data with predictor values drawn from worldwide climate (minimum and maximum temperatures and total June rainfall)(Fick and Hijmans, 2017) and city demographics (Brinkhoff, 2023) (total population and population density). Specifically, covariates for a given city are considered minimum and maximum temperatures and total rainfall for June, total population, and population density.

Dirichlet regression (Maier, 2014) is a technique designed explicitly for relating a compositional response to covariates. With its denomination originating from a namesake probability distribution, the model setup involves multiplicative parameters associated with each predictor, testable for significance and interpretable regarding the context. The results, detailed in Section 4.7, establish that all predictors are statistically significant in determining the response distribution except for rainfall.

While it was fortuitous that precisely *three* proportions were chosen to be studied, this conveniently allowed results to be represented graphically using *ternary plots* (Hamilton and Ferry, 2018). These are designed to represent triplets of percentages that add to 100%, working with axes oriented along the sides of an equilateral triangle so that its barycenter corresponds to value (1/3, 1/3, 1/3). Each vertex represents 100% in exactly one of the components (Figure 5). The vast amount of variability in this type of data becomes conspicuously evident, yet with points tending to gravitate towards the axis that corresponds to 100% *Enterobacter* and 0% *Escherichia* (Figure 5a). Dirichlet regression probes into any structure that may be present in these scattered points. A definite relationship is determined (Section 4.7) with demographic and temperature variables; when quantified in terms of their effect on proportion means, it becomes easier to detect differences between cities (Figure 5b). Analysis corroborates notable variability, with trends towards > 80% *Enterobacter* and < 20% *Escherichia*. Dirichlet regression uncovered underlying structures within the data, revealing statistically significant relationships with demographic and temperature variables. Quantifying their effects on mean abundances facilitated the detection of differences amongst cities. In addition, we posited that the fitted model could be used to simulate random instances of compositional vectors to investigate hypothetical scenarios based on varying predictor values.

**Figure 5.**
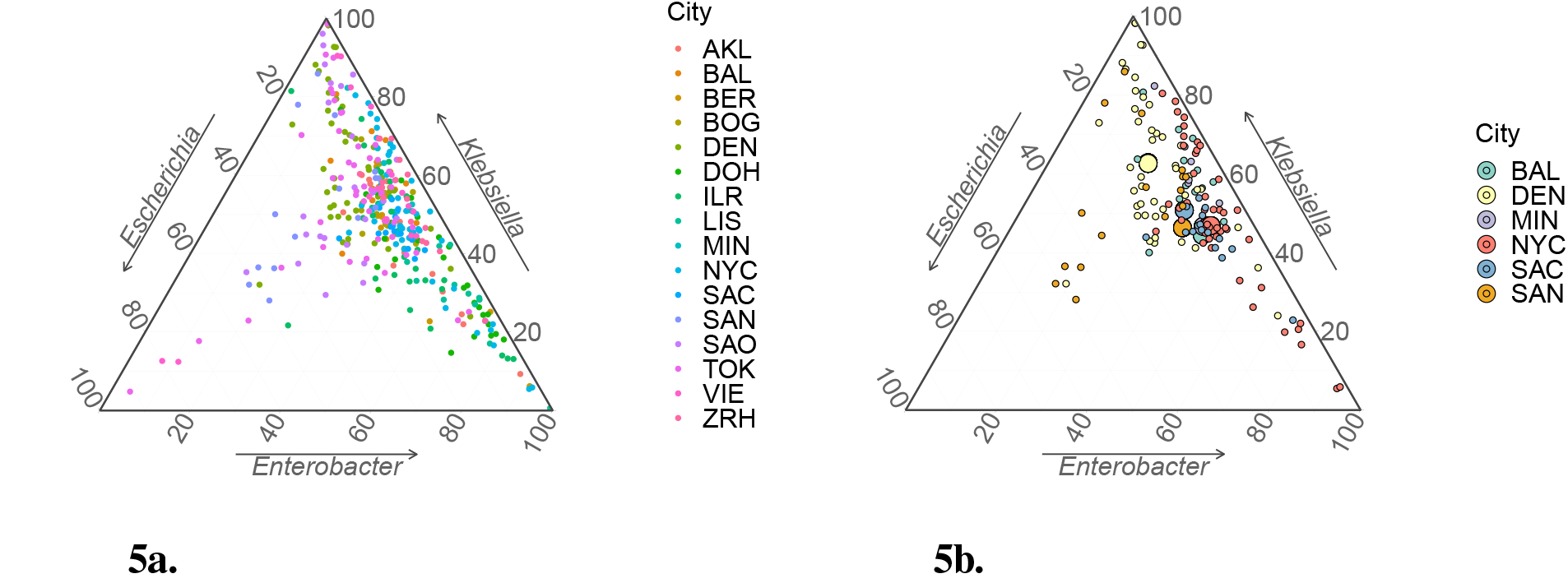
Ternary plots for compositional sample data. **(a)** All 366 data points in database. **(b)** Data points from USA cities only (small circles) with estimated vectors of mean values via Dirichlet regression, (*µ*_1_, *µ*_2_, *µ*_3_), superimposed (large circles). Small circles thus depict raw data, whereas positions of large circles implicitly incorporate climatic and demographic information. The precise nature of these variables’ effects on distributions of relative abundance are elicited from Table 4.

## 3 DISCUSSION

Forensic metagenomics community has tested several variable reduction and classification algorithms as well as proposed new approaches to obtain information from microbiological data. In variable selection, clustering and dimensionality reduction, techniques such as t-SNE (Casimiro-Soriguer et al., 2019; Ryan, 2019), PCA (Walker and Datta, 2019), and UMAP have been employed. Among classification methods Support Vector Machines (Walker and Datta, 2019; Zhu et al., 2019), Random Forests (Walker et al., 2018; Walker and Datta, 2019; Ryan, 2019), and Neural Networks (Zhelyazkova et al., 2021) have been extensively used to identify the city of origin of urban microbiomes. A constant goal has been to identify bacterial fingerprints for the provided cities. To produce fingerprints taxonomic (Walker and Datta, 2019; Ryan, 2019; Danko et al., 2021), functional (Zhu et al., 2019; Casimiro-Soriguer et al., 2019; Danko et al., 2021), and antibiotic resistance(Casimiro-Soriguer et al., 2019; Zhelyazkova et al., 2021; Danko et al., 2021) features have been considered. To construct the fingerprints, organisms that maximize the differences between cities are selected (Walker and Datta, 2019).

**Table 4.**
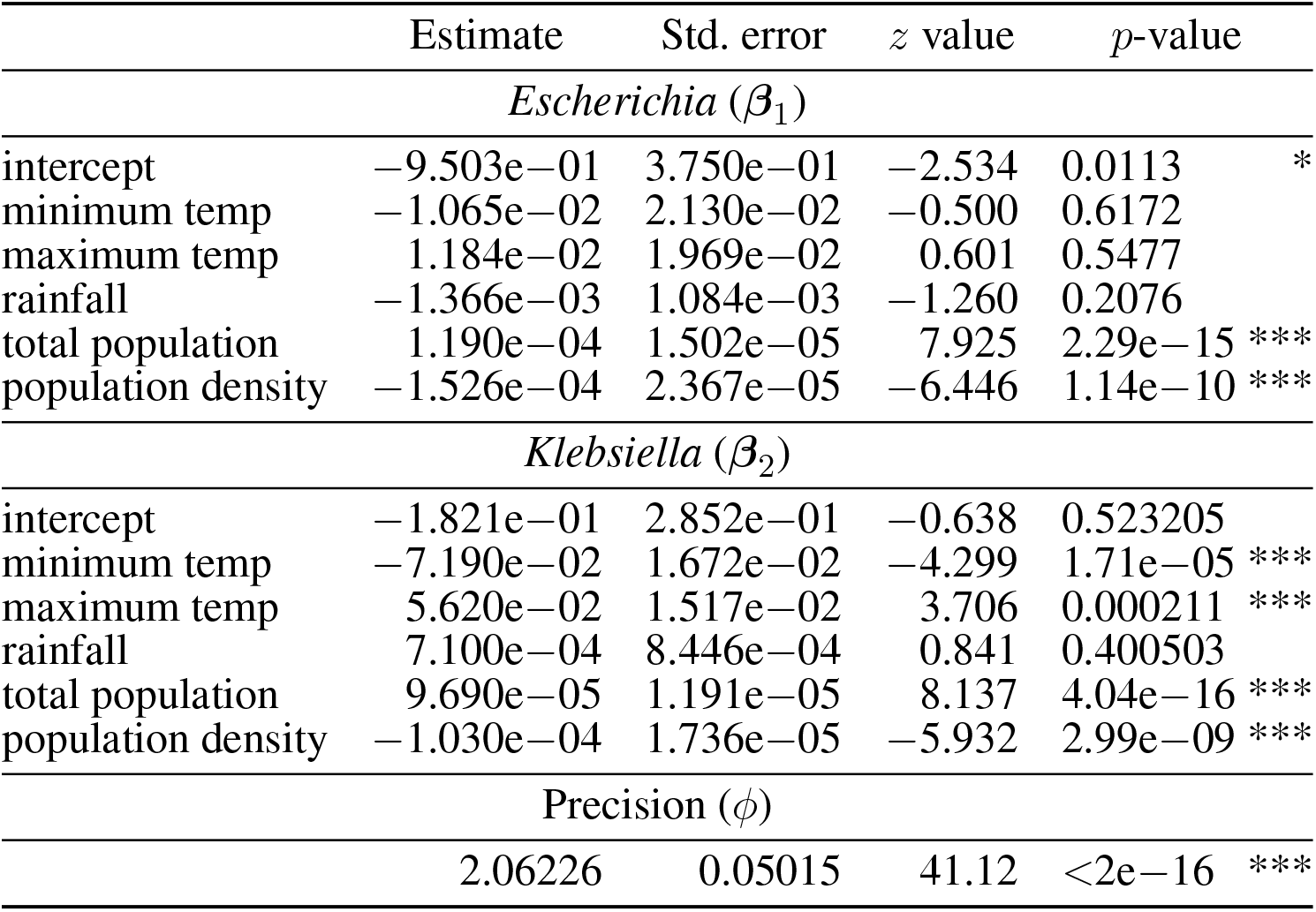
Estimates in the Dirichlet regression model using the parametrization described in the text. Estimates for *Enterobacter* are not required because it is implicitly acting as a reference category. The *z* test statistics and associated *p*-values refer to the test of the hypothesis that the corresponding parameter is zero. Asterisks denote standard R codes for significance: ∗∗∗ = 0, ∗∗= 0.0001, ∗ = 0.01. With *Enterobacter* adopted as the reference, these are examples of how individual estimates are to be interpreted: 1.190e −04 for *Escherichia* under population is highly significant and positive, meaning that an increase in population is associated with a greater mean proportion of *Escherichia* relative to *Enterobacter*; −7.190e−02 for *Klebsiella* under minimum temperature is highly significant and negative, so that a lower minimum temperature is associated with a higher proportion of *Klebsiella*; rainfall is not significant for neither *Escherichia* nor *Klebsiella*.

|  | Estimate | Std. error | *z* value | *p*-value |  |
| --- | --- | --- | --- | --- | --- |
| *Escherichia* ($\beta_1$) | | | | | |
| intercept | -9.503e-01 | 3.750e-01 | -2.534 | 0.0113 | * |
| minimum temp | -1.065e-02 | 2.130e-02 | -0.500 | 0.6172 |  |
| maximum temp | 1.184e-02 | 1.969e-02 | 0.601 | 0.5477 |  |
| rainfall | -1.366e-03 | 1.084e-03 | -1.260 | 0.2076 |  |
| total population | 1.190e-04 | 1.502e-05 | 7.925 | 2.29e-15 | *** |
| population density | -1.526e-04 | 2.367e-05 | -6.446 | 1.14e-10 | *** |
| *Klebsiella* ($\beta_2$) | | | | | |
| intercept | -1.821e-01 | 2.852e-01 | -0.638 | 0.523205 |  |
| minimum temp | -7.190e-02 | 1.672e-02 | -4.299 | 1.71e-05 | *** |
| maximum temp | 5.620e-02 | 1.517e-02 | 3.706 | 0.000211 | *** |
| rainfall | 7.100e-04 | 8.446e-04 | 0.841 | 0.400503 |  |
| total population | 9.690e-05 | 1.191e-05 | 8.137 | 4.04e-16 | *** |
| population density | -1.030e-04 | 1.736e-05 | -5.932 | 2.99e-09 | *** |
| Precision ($\phi$) | | | | | |
|  | 2.06226 | 0.05015 | 41.12 | <2e-16 | *** |

The NB model’s wide applications (Lu et al., 2005) inspired us to adopt methodologies from differential gene expression analyses to select OTUs exhibiting significant count variations across at least two year-cities. Specifically, under the NB model, selected OTUs exhibit greater variability in counts than expected by chance alone. In our work the OTU that were more times differential in pair comparisons between year-cities were *Staphylococcus* (75 times), *Cutibacterium* (54 times), *Stutzerimonas* (46 times), *Bradyrhizobium* (46 times), and *Hydrogenophilus* (40 times), see Supplementary Table S1. Interestingly, *Cutibacterium acnes* and *Bradyrhizobium* sp. BTAI1 were the most relatively abundant species in the global Urban Microbiome calculated in 2021 (Danko et al., 2021), and *Cutibacterium*, is one of the most common bacteria in the skin microbiome. In contrast some species of the genera *Acinetobacter, Pseudomonas* and *Janthinobacter* were selected for being more differential in Walker 2019 study (Walker and Datta, 2019), while *Campylobacter jejuni* and *Staphylococcus argenteus* were found highly predictive in Ryan 2019 work (Ryan, 2019). Discrepancies in differential species are maybe due to the method of variable and selection, and of course to the fact that each year the targeted cities can be different.

Functional profiles achieve lower F1 scores than taxonomic classification which is in agreement with the observation global in the metagenomic map of urban microbiomes (Danko et al., 2021) that functional profiles are more homogeneous across urban samples than taxonomic profiles. Nevertheless this low performance could also be due to the fact that functional categories are more focused on conserved functions. Perhaps functions that differentiate cities are contained in families of specialized functions that are not known yet and in consequence we are not using them as features. Our classification using Mifaser functional annotations achieved a commendable 0.7 accuracy, aligning with the state of the art of the field. This finding is particularly noteworthy compared to existing studies that utilized KEGG annotations, where reported accuracies reached 0.73 using a 10-fold CV(Casimiro-Soriguer et al., 2019). Our results demonstrate a competitive accuracy level, reinforcing the efficacy of our chosen approach and emphasizing the relevance of Mifaser annotations in achieving outcomes comparable to those of widely used databases like KEGG. Our obtained results align with and contribute to the field’s current state of the art. Specifically, through a 10-fold cross-validation, our analysis using Mifaser at specificity level 4 achieved a commendable 70% accuracy. This finding is particularly noteworthy compared to existing studies utilizing KEGG annotations, where reported accuracies reached 73% using a 10-fold cross-validation schema (Casimiro-Soriguer et al., 2019). The slight variation in performance could be attributed to the differences in annotation databases and methodologies employed. Nonetheless, our results demonstrate a competitive accuracy level, reinforcing the efficacy of our chosen approach and emphasizing the relevance of Mifaser annotations in achieving comparable outcomes to widely used databases like KEGG.

Regarding the three bacterial genera of main interest described above as related to city covariates, we found that increased population density (total population) is significantly associated with lower (higher) average proportions of *Escherichia* and *Klebsiella*, relative to *Enterobacter*. This contrasts with Zhang et al. (Zhang et al., 2021b), who proposed considering city-specific metadata such as weather data to improve prediction outcomes, and suggested that environmental factors play a more influential role in microbial composition (Jansson and Hofmockel, 2020) than do demographic variables. Incorporating a broader collection of covariates that included geographical aspects as well, Danko et al. (Danko et al., 2021) established that climate-related components significantly differentiated samples, whereas, admittedly to their surprise, population density did not display a significant effect on taxonomic variation. However, these authors acknowledge a possible masking effect due to correlations between covariates. In any case, albeit these inconsistencies in results may stem from differences in study scope and methods, it is demonstrated that the effects of environmental, demographic and geographic variables on microbial composition merits further, explicit research.

For future studies, it would be valuable to benchmark the potential enhancements in performance through the combination of different taxonomical and functional profiles. Investigating the synergies between various annotation tools or databases could lead us to more robust models and better predictions. Additionally, OTUS selected by different feature selection techniques could be compared and ranked with some score according to how many times different studies identify them as part of the most informative features contributing to the classification profiles of some specific year-city. This approach may not only improve the interpretability of the models but also potentially enhance their predictive performance.

## 4 METHODOLOGY

### 4.1 Data prepocessing

Integrity was checked on the WGS metagenomic paired-end samples by parsing each file with the SeqIO module of BioPython 1.78 (Cock et al., 2009); any sample that failed this test was excluded from downstream analyses. The resulting samples were then adapter and quality trimmed with TrimGalore 0.6.10 (Krueger et al., 2023), discarding reads shorter than 40 base pairs. Afterwards, assemblies were performed at sample- and city-levels using MEGAHIT 1.2.9 (Li et al., 2015).

### 4.2 OTU abundance tables

The taxonomic profiles were predicted with Kraken 2.1.3 (Wood et al., 2019) from read and assembly data at both levels above, using the March 14th, 2023 version of Kraken’s database available as an AWS S3 Bucket (Langmead, 2023). The taxonomic abundance tables were produced in BIOM format (McDonald et al., 2012) with kraken-biom 1.2.0 (Dabdoub, 2016) using sample taxonomy data.

### 4.3 Functional profiles tables

Functional profiles were annotated with two different pipelines: mi-faser 1.60 (Zhu et al., 2017) on the reads and one of EnvGen’s Metagenomics Workshop functional annotation pipelines (Alneberg et al., 2014) on the contigs. The latter, precisely, consists in the following: first, the assemblies are annotated with Prokka 1.14.6 (Seemann, 2014b), modified to skip the execution of tbl2asn, which was found to be extremely slow when working with large metagenomic assemblies; and second, using the E. C. numbers identified by Prokka and the MetaCyc Database (Caspi et al., 2019) as reference, we use MinPath 1.6 (Ye and Doak, 2009) to obtain the minimum set of MetaCyc pathways, which is then hierarchized into eight functional levels.

### 4.4 Variable Selection

To select variables by addressing the zero-inflated phenomenon, we use four regression models for count data 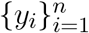. The fitted models consider the covariates ***x***_*i*_ and ***z***_*i*_ to be the categorical variables that correspond to the location from which the samples originate. The statistical fits are implemented with the R (version 4.3.0) packages-functions: stats-`glm`, MASS-`glm.nb` and pscl-`zeroinfl`. These regression models, consider as offset the logarithms of the total read, {log(*N*_*i*_)}, and are:

1. Poisson (P),

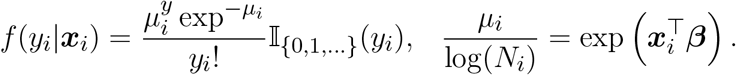
2. Negative binomial (NB),

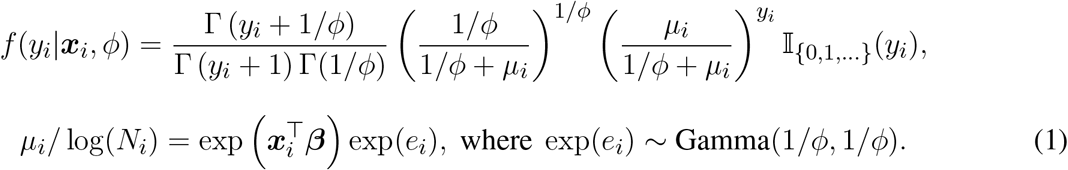
3. Zero inflated Poisson (ZIP),

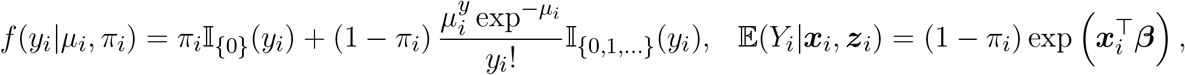 Where the probability of *y*_*i*_ being a zero count is modeled in the regression model with logit function and covariates ***z***_*i*_:

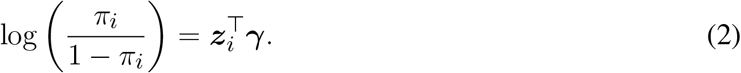
4. Zero inflated negative binomial (ZINB).

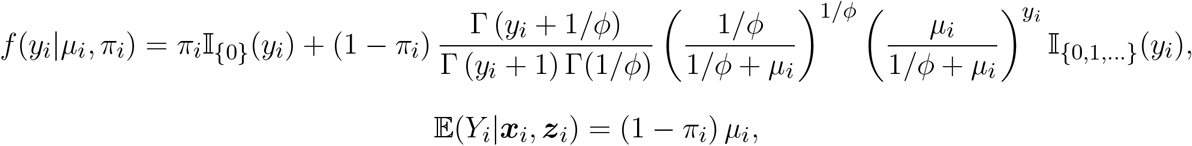

with *µ*_*i*_*/* log(*N*_*i*_), *e*_*i*_ and *π*_*i*_ as in (1) and (2).

This procedure is performed for each of the taxonomic levels previously presented and for each pair of year-cities. However, the variable selection considers all the computed *p*-values resulting from the analysis at each taxonomic level. We only consider pairs of categories where the OTU count is greater than zero for both. This is to avoid numerical errors.

To control the FDR we used the Benjamini-Hochberg procedure (Benjamini and Hochberg, 1995) implemented in R with the command `p.adjust`. Its procedure for variable selections is

1. Sort the *p*-values in ascending order and assigning a rank or position *p*_(1)_ ≤ *p*_(2)_ ≤ · · · ≤ *p*_(*k*)_.
2. Compute the adjusted *p*-value for using the formula:

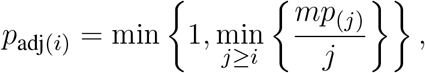

where *m* represents the total number of hypotheses being tested, the purpose of taking the minimum ratio is to find the smallest value that controls the FDR while considering all hypotheses ranked at or above the current *p*-value. Taking the minimum ensures that the adjusted *p*-value is conservative and provides a corrected measure of statistical significance that accounts for multiple tests.

Typically, for a given test level *α*, we find the largest *j* such that *p*_adj(*j*)_ ≤ *α* and reject the null hypothesis (i.e., declare discoveries) for those variables associated to the first *j* ranked *p*-values. However, for identifying the OTUs that can be more successful in differentiating cities (year-cities), we obtained all the *p*-values for each pair of year-cities and selected those OTUs with the lowest *p*-values. The objective was to obtain a reduced list of OTUs that can perform well for the forensic challenge of city classification.

For each model (P, NB, ZIP, and ZINB) we selected the 5 OTUs with the lowest recorded *p*-values. Table 3 presents the resulting number of OTUs under each model, and in parenthesis, we report the percentage of zero counts for the obtained OTUs for all pairs of cities. The resulting OTUs under “Model selection” are those from the best model (P, NB, ZIP or ZINB) that for each pair of cities, has the smallest AIC.

### 4.5 Prediction Models using OTUs selection

Code for city prediction uses Python 3.10, with libraries scikit-learn 1.0.2, pandas 1.5.3, and matplotlib 3.7.1. To establish a robust and consistent prediction process, a 5-fold cross-validation scheme was implemented (Figure 3c). Within this framework, a stratified procedure was devised to partition the initial dataset into five distinct groups. This approach ensured that city proportions remained consistent across all groups. While training, each city was considered as a different class for samples from different years.

The 5-fold process divided the data into one validation set and four training sets per iteration to ensure a thorough evaluation. Subsequently, a variable selection process was conducted exclusively on the training set using the NB method, chosen for its robust performance in preliminary tests. This procedure aimed to guarantee that the selected variables contained no information from the validation set. Following the variable selection process, different selections were made for each fold due to the varying information available in the distinct training sets. The resulting variables from the training set were then utilized for the subsequent classification process.

The initial stage of the classification process involved taking the reduced set of variables from the training set and subjecting it to standardization and normalization procedures (using python’s sklearn libraries) to prepare the data for classification algorithms. Extensive testing identified the quantile transformer (normalizing by OTUs instead of cities) with z-score as the most effective algorithm.

The subsequent phase of the classification process encompassed the application of three potential algorithms: MLP, KNN, and SVC, all from the sklearn libraries. To ensure reproducibility, all algorithms employed a fixed random seed.

The specific configurations utilized for each algorithm were 200 neurons in a single layer for MLP, 23 neighbors for KNN, and a linear kernel with 2 degrees for SVC. Although several other hyperparameters were assessed, including some random forest classifier models, they exhibited subpar performance.

For each cross-validation fold, the Macro F1 score was computed for each of the algorithms mentioned above, understanding that before this process, all the years for a given city were merged into a single class representing that city.

### 4.6 Classification models with functional profiles

To determine the most effective parameter configurations for the models, we employed Grid Search, a systematic approach exploring a range of hyperparameters, see Supplementary material Table S3. For the SVC, the optimal configuration involved a linear kernel, chosen after thorough evaluation during Grid Search. The MLP model achieved optimal results with specific parameters identified through Grid Search. These included a Tanh activation function, automatic batch size adjustment, disabled early stopping, a single hidden layer of 100 neurons, adaptive learning rate, a maximum of 3000 training iterations, and the Stochastic Gradient Descent (SGD) solver.

Similarly, for the KNN Classifier, Grid Search determined that configuring the model with two neighbors for prediction and a distance-based weighting scheme yielded the most accurate predictions.

We employed a 5-fold cross-validation strategy to assess our models’ robustness and generalization. This involved partitioning the dataset into five subsets, training the models on four subsets, and evaluating their performance on the remaining subset. The average performance across these folds provided a reliable estimate of the overall effectiveness of the models.

### 4.7 Dirichlet regression

A Dirichlet distribution for a random compositional triplet ***Y*** = (*Y*_1_, *Y*_2_, *Y*_3_) is described by three parameters *µ*_1_, *µ*_2_, *µ*_3_ in (0, 1) representing the means of each entry (E(*Y*_*c*_) = *µ*_*c*_), plus a precision parameter, *ϕ >* 0, that controls variability around the means (Maier, 2014). The restriction 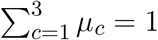 is enforced. Set *X*_0_ = 1 to allow for an intercept term. If *C* predictors are aggregated and denoted by ***X*** = (*X*_0_, *X*_1_, …, *X*_*C*_) and a set of coefficients by ***β*** = (*β*_0_, *β*_1_, …, *β*_*C*_)^T^, we use vector notation to write 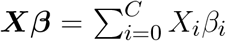 Dirichlet regression postulates that mean parameters depend on covariates by setting

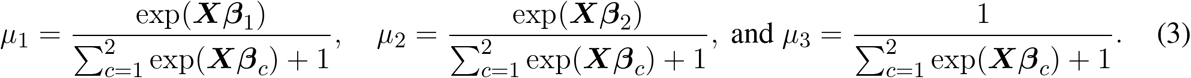

Regression parameters are ***β***_1_ and ***β***_2_, one vector for each of the first two components, plus the precision parameter, *ϕ*. A third beta parameter, ***β***_3_ say, is neither involved nor required because the restriction 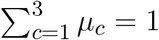 determines the third mean in terms of the other two. This parametrization implicitly signifies that the third component is viewed as a reference category in terms of ratios between means. More precisely, interpretation of ***β***_1_ and ***β***_2_ follows from noting that *µ*_1_*/µ*_3_ = exp(***Xβ***_1_) and *µ*_2_*/µ*_3_ = exp(***Xβ***_2_). In fact, if a different category were to be chosen as the reference, the same estimated values of *µ*_1_, *µ*_2_ and *µ*_3_ would result; the reference category simply establishes one baseline to compare the other two against.

In addition to the means in (3) as specific properties of the Dirichlet distribution, we may also state

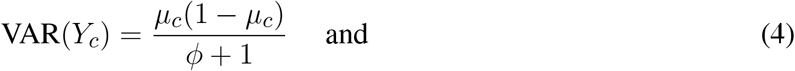

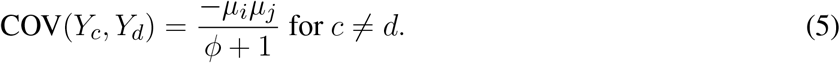

The reason for referring to *ϕ* as a *precision* parameter becomes clear from (4), because a larger value of *ϕ* results in smaller variances of all proportions in the compositional vector ***Y***. Negative covariance in (5) is also expected since a larger proportion in any given entry ensues at the expense of smaller proportions in the other two.

Package `DirichletReg` in R (Maier, 2021) addresses Dirichlet regression and implements model fitting by numerical methods for maximum likelihood. Estimated parameters (Table 4) grant a Dirichlet density to describe the distribution of compositional triplets for a given value of ***X*** pertaining to a particular city. The regression structure exploited across cities means that assessments can be made for hypothetical new values of ***X***. For example, the effect on bacterial composition if maximum temperature were to increase by 1°C or if population density increases by 2%. Estimated values of *µ*_1_, *µ*_2_, *µ*_3_ for given values of ***X*** can be readily obtained by plugging-in estimated coefficients shown in Table 4 into equations (3) (as has been done in Figure 5b). Likewise, estimated variances are obtained by plugging into (4).

Another use for regression models is the simulation of specified scenarios. Once model parameters have been estimated based on data, clouds of simulated (pseudo) data points can be obtained for understanding the complexion of variability or for comparing probabilistic distributions at different levels of ***X***. To enable this, R package `DirichletReg` provides function `rdirichlet` to simulate random instances of compositional vectors for a specified set of parameters.

## AUTHOR CONTRIBUTIONS

AP and RPE contribute to data prepossessing. SGF desing and supervised the data preprocessing pipeline. INM, LLRR and MECB contributed to variable selection. DSQ, EB contributed to prediction using abundance tables. MVRL contributed to functional annotation. RPE contributed to prediction models using functional annotation. VMS and SGF contributed to classification models. HCP and MNS contributed to Dirichlet regression models. AP, RPE, LLRR, DSQ, MNS, HCP and NS wrote the manuscript. LLRR, MNS, MNS, NS and HCP revised the article.

### FUNDING

NSM was supported by CONAHCYT grant 320237. This work was supported by UNAM Posdoctoral Program (POSDOC) at Centro de Ciencias Matemáticas. We thank the financial support of DGAPA grant PAPIIT IN101423.

## ACKNOWLEDGMENTS

We thank CCM, CIMAT, and UNAM:IryA, and UNAM-Huawei-Alianza for the allowance of computer servers and for amazing technical support.

## DATA AVAILABILITY STATEMENT

The datasets [GENERATED/ANALYZED] for this study can be found in the [ccm-bioinfo/cambda2023].

