## Supplementary Material for "CAMDA 2023: finding patterns in urban microbiomes"

### 1 SUPPLEMENTARY DATA

#### 1.1 Most differential OTUs

Table S1: We deemed an OTU to be **most differential** if it is differential for more than 10 city comparisons; i.e. if it is selected as differential for at least 11 pairs of cities. With this definition we obtained the following table, where we present the ID of the identified OTUs, the number of times it was identified as differential, and its taxonomic classification (up to Genera).

| OTU | Comparisons | Kingdom | Phylum | Class | Order | Family | Genus |
|---|---|---|---|---|---|---|---|
| 1280 | 75 | Bacteria | Bacillota | Bacilli | Bacillales | Staphylococcaceae | Staphylococcus |
| 1747 | 54 | Bacteria | Actinomycetota | Actinomycetes | Propionibacteriales | Propionibacteriaceae | Cutibacterium |
| 316 | 46 | Bacteria | Pseudomonadota | Gammaproteobacteria | Pseudomonadales | Pseudomonadaceae | Stutzerimonas |
| 288000 | 46 | Bacteria | Pseudomonadota | Alphaproteobacteria | Hyphomicrobiales | Nitrobacteraceae | Bradyrhizobium |
| 297 | 40 | Bacteria | Pseudomonadota | Hydrogenophilia | Hydrogenophilales | Hydrogenophilaceae | Hydrogenophilus |
| 1980928 | 33 | Viruses | Uroviricota | Caudoviricetes | Herelleviridae | Sepunavirus | |
| 2596915 | 27 | Bacteria | Bacteroidota | Cytophagia | Cytophagales | Hymenobacteraceae | Hymenobacter |
| 1458307 | 26 | Bacteria | Pseudomonadota | Alphaproteobacteria | Rhodobacterales | Roseobacteraceae | Octadecabacter |
| 1528099 | 25 | Bacteria | Actinomycetota | Actinomycetes | Mycobacteriales | Lawsonellaceae | Lawsonella |
| 565 | 24 | Bacteria | Pseudomonadota | Gammaproteobacteria | Enterobacterales | Enterobacteriaceae | Atlantibacter |
| 1982251 | 19 | Viruses | Uroviricota | Caudoviricetes | Pahexavirus | | |
| 29370 | 16 | Bacteria | Bacillota | Clostridia | Eubacteriales | Lachnospiraceae | Lacrimispora |
| 225148 | 16 | Bacteria | Chlamydiota | Chlamydiia | Parachlamydiales | Rhabdochlamydiaceae | Candidatus Rhabdochlamydia |
| 2713640 | 16 | Viruses | Artverviricota | Revtraviricetes | Ortervirales | Caulimoviridae | Cavemovirus |
| 49319 | 15 | Bacteria | Actinomycetota | Rubrobacteria | Rubrobacterales | Rubrobacteraceae | Rubrobacter |
| 546871 | 15 | Bacteria | Actinomycetota | Actinomycetes | Propionibacteriales | Nocardioidaceae | Friedmanniella |
| 712435 | 14 | Bacteria | Bacteroidota | Bacteroidia | Bacteroidales | Porphyromonadaceae | Porphyromonas |
| 22 | 12 | Bacteria | Pseudomonadota | Gammaproteobacteria | Alteromonadales | Shewanellaceae | Shewanella |
| 566 | 12 | Bacteria | Pseudomonadota | Gammaproteobacteria | Enterobacterales | Enterobacteriaceae | Pseudescherichia |
| 38304 | 12 | Bacteria | Actinomycetota | Actinomycetes | Mycobacteriales | Corynebacteriaceae | Corynebacterium |
| 165179 | 11 | Bacteria | Bacteroidota | Bacteroidia | Bacteroidales | Prevotellaceae | Prevotella |
| 262209 | 11 | Bacteria | Actinomycetota | Actinomycetes | Micrococcales | Intrasporangiaceae | Janibacter |
| 2086471 | 11 | Bacteria | Bacteroidota | Cytophagia | Cytophagales | Hymenobacteraceae | Adhaeribacter |
| 2939494 | 11 | Bacteria | Bacillota | Bacilli | Bacillales | Planococcaceae | Rummeliibacillus |

### 1.2 Classification models with abundance tables

| abundance | Kingdom | Phylum | Class | Order | Family | Genus | Fold presence |
|---|---|---|---|---|---|---|---|
| 855575815 | Bacteria | Pseudomonadota | NA | NA | NA | NA | 1 1 1 1 1 |
| 618757355 | Bacteria | Pseudomonadota | Gammaproteobacteria | NA | NA | NA | 1 1 1 1 1 |
| 227223542 | Bacteria | Pseudomonadota | Gammaproteobacteria | Pseudomonadales | Pseudomonadaceae | Stutzerimonas | 1 1 1 1 1 |
| 184708059 | Bacteria | Bacillota | NA | NA | NA | NA | 1 1 1 1 1 |
| 157476883 | Bacteria | Pseudomonadota | Alphaproteobacteria | NA | NA | NA | 1 1 0 0 0 |
| 148311860 | Bacteria | Bacillota | Bacilli | NA | NA | NA | 1 1 1 1 1 |
| 146397481 | Bacteria | Pseudomonadota | Gammaproteobacteria | Enterobacterales | NA | NA | 1 1 1 1 1 |
| 132192372 | Bacteria | Pseudomonadota | Gammaproteobacteria | Moraxellales | NA | NA | 1 1 1 1 1 |
| 132192372 | Bacteria | Pseudomonadota | Gammaproteobacteria | Moraxellales | Moraxellaceae | NA | 1 1 1 1 1 |
| 126935116 | Bacteria | Bacillota | Bacilli | Bacillales | NA | NA | 1 1 1 1 1 |
| 117201018 | Bacteria | Pseudomonadota | Gammaproteobacteria | Moraxellales | Moraxellaceae | Acinetobacter | 1 1 0 0 1 |
| 100087449 | Bacteria | Actinomycetota | Actinomycetes | Propionibacteriales | NA | NA | 1 1 1 1 1 |
| 79516422 | Bacteria | Actinomycetota | Actinomycetes | Propionibacteriales | Propionibacteriaceae | NA | 1 1 1 1 1 |
| 72833080 | Bacteria | Actinomycetota | Actinomycetes | Propionibacteriales | Propionibacteriaceae | Cutibacterium | 1 1 1 1 1 |
| 64434208 | Bacteria | Pseudomonadota | Alphaproteobacteria | Hyphomicrobiales | NA | NA | 1 1 0 0 1 |
| 41773446 | Bacteria | Bacillota | Bacilli | Bacillales | Bacillaceae | Lysinibacillus | 1 1 1 1 1 |
| 39764946 | Bacteria | Actinomycetota | Actinomycetes | Micrococcales | Micrococcaceae | NA | 1 1 1 1 1 |
| 38340331 | Bacteria | Bacillota | Bacilli | Bacillales | Staphylococcaceae | NA | 1 1 1 1 1 |
| 37537514 | Bacteria | Pseudomonadota | Gammaproteobacteria | Enterobacterales | Erwiniaceae | NA | 1 1 1 1 1 |
| 37527310 | Bacteria | Bacillota | Bacilli | Bacillales | Staphylococcaceae | Staphylococcus | 1 1 1 1 1 |
| 36421196 | Bacteria | Pseudomonadota | Alphaproteobacteria | Sphingomonadales | NA | NA | 1 1 0 1 0 |
| 36046769 | Bacteria | Pseudomonadota | Gammaproteobacteria | Xanthomonadales | NA | NA | 1 0 1 1 0 |
| 35017217 | Bacteria | Pseudomonadota | Gammaproteobacteria | Xanthomonadales | Xanthomonadaceae | NA | 1 0 1 1 0 |
| 30275392 | Bacteria | Pseudomonadota | Gammaproteobacteria | Enterobacterales | Erwiniaceae | Pantoea | 1 1 0 1 1 |
| 29382471 | Bacteria | Pseudomonadota | Alphaproteobacteria | Sphingomonadales | Sphingomonadaceae | NA | 1 1 0 1 1 |
| 27966060 | Bacteria | Pseudomonadota | Alphaproteobacteria | Hyphomicrobiales | Nitrobacteraceae | NA | 1 1 1 1 1 |
| 26695213 | Bacteria | Pseudomonadota | Alphaproteobacteria | Hyphomicrobiales | Nitrobacteraceae | Bradyrhizobium | 1 1 1 1 1 |
| 24460060 | Bacteria | Actinomycetota | Actinomycetes | Mycobacteriales | Corynebacteriaceae | NA | 1 1 1 1 1 |
| 24460060 | Bacteria | Actinomycetota | Actinomycetes | Mycobacteriales | Corynebacteriaceae | Corynebacterium | 1 1 1 1 1 |
| 24433767 | Bacteria | Pseudomonadota | Betaproteobacteria | Burkholderiales | Oxalobacteraceae | NA | 1 0 1 1 1 |
| 23777536 | Bacteria | Pseudomonadota | Gammaproteobacteria | Xanthomonadales | Xanthomonadaceae | Stenotrophomonas | 1 0 1 1 0 |
| 21376744 | Bacteria | Bacillota | Bacilli | Lactobacillales | NA | NA | 1 0 0 0 0 |
| 18567411 | Bacteria | Pseudomonadota | Gammaproteobacteria | Enterobacterales | Enterobacteriaceae | Leclercia | 1 1 1 1 1 |

| | | | | | | | |
|---|---|---|---|---|---|---|---|
| 18279758 | Bacteria | Pseudomonadota | Alphaproteobacteria | Sphingomonadales | Sphingomonadaceae | Sphingomonas | 1 1 1 1 1 |
| 18096618 | Bacteria | Pseudomonadota | Alphaproteobacteria | Rhodobacterales | Paracoccaceae | Paracoccus | 1 1 1 1 1 |
| 15830629 | Bacteria | Actinomycetota | Actinomycetes | Micrococcales | Micrococcaceae | Micrococcus | 1 0 0 1 1 |
| 15653805 | Bacteria | Bacillota | Clostridia | Eubacteriales | Lachnospiraceae | NA | 1 0 0 0 0 |
| 12376123 | Bacteria | Actinomycetota | Actinomycetes | Micrococcales | Micrococcaceae | Kocuria | 1 1 1 1 1 |
| 12133394 | Bacteria | Bacteroidota | Flavobacteriia | Flavobacteriales | Weeksellaceae | NA | 1 1 1 1 1 |
| 11832944 | Bacteria | Pseudomonadota | Alphaproteobacteria | Hyphomicrobiales | Methylobacteriaceae | NA | 1 1 1 1 1 |
| 11778825 | Bacteria | Pseudomonadota | Gammaproteobacteria | Moraxellales | Moraxellaceae | Moraxella | 1 1 1 1 1 |
| 9917037 | Bacteria | Bacteroidota | Cytophagia | Cytophagales | NA | NA | 1 1 1 0 1 |
| 9917037 | Bacteria | Bacteroidota | Cytophagia | NA | NA | NA | 1 1 1 1 1 |
| 9430242 | Bacteria | Pseudomonadota | Alphaproteobacteria | Caulobacterales | Caulobacteraceae | NA | 1 0 0 0 0 |
| 8834074 | Bacteria | Pseudomonadota | Alphaproteobacteria | Hyphomicrobiales | Methylobacteriaceae | Methylobacterium | 1 1 1 1 1 |
| 8507543 | Bacteria | Bacteroidota | Cytophagia | Cytophagales | Hymenobacteraceae | NA | 1 1 1 1 1 |
| 8023338 | Bacteria | Bacteroidota | Cytophagia | Cytophagales | Hymenobacteraceae | Hymenobacter | 1 1 1 1 1 |
| 7005042 | Bacteria | Actinomycetota | Actinomycetes | Micrococcales | Intrasporangiaceae | NA | 1 1 1 1 1 |
| 5268123 | Bacteria | Bacteroidota | Flavobacteriia | Flavobacteriales | Weeksellaceae | Chryseobacterium | 1 1 1 1 1 |
| 4739621 | Bacteria | Pseudomonadota | Betaproteobacteria | Burkholderiales | Burkholderiaceae | Cupriavidus | 1 1 1 1 1 |
| 4433102 | Bacteria | Actinomycetota | Actinomycetes | Micrococcales | Intrasporangiaceae | Janibacter | 1 1 1 1 1 |
| 4407862 | Bacteria | Bacteroidota | Bacteroidia | NA | NA | NA | 1 1 1 1 1 |
| 4354102 | Bacteria | Bacteroidota | Bacteroidia | Bacteroidales | NA | NA | 1 1 0 1 1 |
| 4249631 | Bacteria | Actinomycetota | Actinomycetes | Geodermatophilales | Geodermatophilaceae | Modestobacter | 1 1 0 0 1 |
| 4196877 | Bacteria | Pseudomonadota | Alphaproteobacteria | Rhodospirillales | Acetobacteraceae | Roseomonas | 1 0 0 0 0 |
| 3683335 | Bacteria | Actinomycetota | Actinomycetes | Micrococcales | Micrococcaceae | Rothia | 1 1 1 1 1 |
| 3651044 | Bacteria | Pseudomonadota | Betaproteobacteria | Burkholderiales | Comamonadaceae | Variovorax | 1 0 1 0 0 |
| 3364490 | Bacteria | Actinomycetota | Actinomycetes | Micrococcales | Ornithinimicrobiaceae | NA | 1 0 0 0 1 |
| 2927152 | Bacteria | Pseudomonadota | Gammaproteobacteria | Moraxellales | Moraxellaceae | Psychrobacter | 1 1 1 1 1 |
| 2858266 | Bacteria | Actinomycetota | Actinomycetes | Propionibacteriales | Propionibacteriaceae | Microlunatus | 1 1 1 1 1 |
| 2789044 | Bacteria | Pseudomonadota | Gammaproteobacteria | Enterobacterales | Erwiniaceae | Mixta | 1 1 1 1 1 |
| 2697058 | Bacteria | Pseudomonadota | Gammaproteobacteria | Alteromonadales | NA | NA | 1 1 1 1 1 |
| 2581926 | Bacteria | Actinomycetota | Actinomycetes | Micrococcales | Dermacoccaceae | Dermacoccus | 1 1 1 1 1 |
| 2408915 | Bacteria | Pseudomonadota | Alphaproteobacteria | Hyphomicrobiales | Phyllobacteriaceae | Mesorhizobium | 1 0 0 0 0 |
| 2119548 | Bacteria | Actinomycetota | Actinomycetes | Micrococcales | Ornithinimicrobiaceae | Ornithinimicrobium | 1 1 0 0 1 |
| 2108076 | Bacteria | Pseudomonadota | Alphaproteobacteria | Hyphomicrobiales | Methylobacteriaceae | Methylorubrum | 1 1 1 1 1 |
| 2106717 | Bacteria | Pseudomonadota | Betaproteobacteria | Burkholderiales | Oxalobacteraceae | Pseudoduganella | 1 1 1 0 0 |
| 2099886 | Bacteria | Actinomycetota | Actinomycetes | Micrococcales | Brevibacteriaceae | NA | 1 1 1 1 0 |
| 2099886 | Bacteria | Actinomycetota | Actinomycetes | Micrococcales | Brevibacteriaceae | Brevibacterium | 1 1 1 1 0 |

| | | | | | | | |
|---|---|---|---|---|---|---|---|
| 2044244 | Bacteria | Pseudomonadota | Gammaproteobacteria | Enterobacterales | Enterobacteriaceae | Escherichia | 1 0 0 0 0 |
| 2036773 | Bacteria | Pseudomonadota | Betaproteobacteria | Burkholderiales | Comamonadaceae | Comamonas | 1 1 0 0 0 |
| 1826196 | Bacteria | Bacillota | Tissierellia | NA | NA | NA | 1 0 0 0 0 |
| 1795720 | Bacteria | Pseudomonadota | Gammaproteobacteria | Alteromonadales | Shewanellaceae | NA | 1 1 1 1 1 |
| 1785818 | Bacteria | Pseudomonadota | Gammaproteobacteria | Alteromonadales | Shewanellaceae | Shewanella | 1 1 1 1 1 |
| 1742405 | Bacteria | Pseudomonadota | Gammaproteobacteria | Enterobacterales | Enterobacteriaceae | Salmonella | 1 1 0 1 1 |
| 1673835 | Bacteria | Bacillota | Bacilli | Lactobacillales | Lactobacillaceae | Leuconostoc | 1 0 0 0 1 |
| 1662215 | Bacteria | Pseudomonadota | Alphaproteobacteria | Caulobacterales | Caulobacteraceae | Caulobacter | 1 1 0 0 0 |
| 1590662 | Bacteria | Bacillota | Tissierellia | Tissierellales | NA | NA | 1 0 0 0 0 |
| 1568927 | Bacteria | Fusobacteriota | Fusobacteriia | Fusobacteriales | NA | NA | 1 1 1 1 1 |
| 1568927 | Bacteria | Fusobacteriota | NA | NA | NA | NA | 1 1 1 1 1 |
| 1568927 | Bacteria | Fusobacteriota | Fusobacteriia | NA | NA | NA | 1 1 1 1 1 |
| 1566693 | Bacteria | Bacillota | Negativicutes | Veillonellales | NA | NA | 1 1 0 0 0 |
| 1566693 | Bacteria | Bacillota | Negativicutes | Veillonellales | Veillonellaceae | NA | 1 1 0 0 0 |
| 1448437 | Bacteria | Pseudomonadota | Gammaproteobacteria | Xanthomonadales | Xanthomonadaceae | Pseudoxanthomonas | 1 1 0 0 1 |
| 1365640 | Bacteria | Bacillota | Bacilli | Lactobacillales | Aerococcaceae | Aerococcus | 1 0 1 1 0 |
| 1335044 | Bacteria | Actinomycetota | Rubrobacteria | Rubrobacterales | NA | NA | 1 0 1 0 0 |
| 1335044 | Bacteria | Actinomycetota | Rubrobacteria | NA | NA | NA | 1 0 1 0 1 |
| 1281711 | Bacteria | Bacteroidota | Flavobacteriia | Flavobacteriales | Weeksellaceae | Epilithonimonas | 1 1 1 1 1 |
| 1164329 | Bacteria | Bacillota | Bacilli | Bacillales | | Exiguobacterium | 1 0 1 0 0 |
| 1142651 | Bacteria | Actinomycetota | Actinomycetes | Micrococcales | Microbacteriaceae | Rathayibacter | 1 1 0 1 1 |
| 1056327 | Bacteria | Bacillota | Bacilli | Lactobacillales | Enterococcaceae | NA | 1 1 0 1 0 |
| 1015128 | Bacteria | Pseudomonadota | Betaproteobacteria | Burkholderiales | Burkholderiaceae | Ralstonia | 1 0 1 1 1 |
| 923404 | Bacteria | Fusobacteriota | Fusobacteriia | Fusobacteriales | Fusobacteriaceae | NA | 1 0 0 0 0 |
| 918726 | Bacteria | Bacillota | Bacilli | Lactobacillales | Enterococcaceae | Enterococcus | 1 1 0 1 1 |
| 897256 | Bacteria | Fusobacteriota | Fusobacteriia | Fusobacteriales | Fusobacteriaceae | Fusobacterium | 1 0 0 0 0 |
| 850434 | Bacteria | Bacteroidota | Chitinophagia | Chitinophagales | NA | NA | 1 0 0 0 0 |
| 850434 | Bacteria | Bacteroidota | Chitinophagia | Chitinophagales | Chitinophagaceae | NA | 1 0 0 0 0 |
| 850434 | Bacteria | Bacteroidota | Chitinophagia | NA | NA | NA | 1 0 0 0 0 |
| 789318 | Bacteria | Bacillota | Clostridia | Eubacteriales | Peptostreptococcaceae | NA | 1 0 0 0 0 |
| 649206 | Bacteria | Pseudomonadota | Alphaproteobacteria | Hyphomicrobiales | Nitrobacteraceae | Rhodopseudomonas | 1 0 0 0 0 |
| 614260 | Bacteria | Pseudomonadota | Alphaproteobacteria | Hyphomicrobiales | Methylobacteriaceae | Microvirga | 1 1 0 0 1 |
| 611115 | Bacteria | Bacillota | Bacilli | Lactobacillales | Lactobacillaceae | Limosilactobacillus | 1 0 0 0 0 |
| 606165 | Bacteria | Pseudomonadota | Alphaproteobacteria | Hyphomicrobiales | Rhizobiaceae | Shinella | 1 0 0 0 0 |
| 568724 | Bacteria | Bacteroidota | Flavobacteriia | Flavobacteriales | Flavobacteriaceae | Capnocytophaga | 1 0 0 0 0 |
| 561983 | Bacteria | Actinomycetota | Actinomycetes | Micrococcales | Microbacteriaceae | Clavibacter | 1 1 0 0 1 |

| | | | | | | | |
|---|---|---|---|---|---|---|---|
| 410778 | Bacteria | Campylobacterota | Epsilonproteobacteria | Campylobacterales | Arcobacteraceae | NA | 1 1 1 1 1 |
| 409880 | Bacteria | Bacillota | Bacilli | Bacillales | Bacillaceae | Peribacillus | 1 1 1 1 0 |
| 391908 | Bacteria | Pseudomonadota | Gammaproteobacteria | Enterobacterales | Yersiniaceae | Rahnella | 1 1 1 1 1 |
| 385880 | Bacteria | Bacillota | Bacilli | Lactobacillales | Lactobacillaceae | Latilactobacillus | 1 1 1 1 1 |
| 344059 | Bacteria | Bacteroidota | Bacteroidia | Bacteroidales | Porphyromonadaceae | NA | 1 1 0 1 1 |
| 344059 | Bacteria | Bacteroidota | Bacteroidia | Bacteroidales | Porphyromonadaceae | Porphyromonas | 1 1 0 1 1 |
| 325807 | Bacteria | Pseudomonadota | Alphaproteobacteria | Sphingomonadales | Erythrobacteraceae | Porphyrobacter | 1 0 0 0 0 |
| 246520 | Bacteria | Pseudomonadota | Gammaproteobacteria | Enterobacterales | Morganellaceae | Proteus | 1 1 1 1 1 |
| 229052 | Bacteria | Bacteroidota | Cytophagia | Cytophagales | Spirosomaceae | Dyadobacter | 1 0 0 0 0 |
| 216431 | Bacteria | Pseudomonadota | Betaproteobacteria | Burkholderiales | Oxalobacteraceae | Collimonas | 1 0 1 0 0 |
| 173556 | Bacteria | Bacillota | Bacilli | Bacillales | Staphylococcaceae | Mammaliicoccus | 1 1 1 1 1 |
| 119209 | Bacteria | Pseudomonadota | Gammaproteobacteria | Enterobacterales | Yersiniaceae | Rouxiella | 1 0 0 0 0 |
| 90232 | Archaea | Euryarchaeota | Halobacteria | Halobacteriales | Halobacteriaceae | Halobacterium | 1 1 0 0 0 |
| 23597355 | Bacteria | Pseudomonadota | Gammaproteobacteria | Enterobacterales | Enterobacteriaceae | Enterobacter | 0 1 0 0 0 |
| 21805113 | Bacteria | Pseudomonadota | Alphaproteobacteria | Rhodospirillales | NA | NA | 0 1 0 0 0 |
| 11296980 | Bacteria | Bacillota | Bacilli | Lactobacillales | Streptococcaceae | NA | 0 1 0 0 0 |
| 9165060 | Bacteria | Bacillota | Bacilli | Lactobacillales | Streptococcaceae | Streptococcus | 0 1 0 0 0 |
| 5895356 | Bacteria | Pseudomonadota | Alphaproteobacteria | Rhodospirillales | Acetobacteraceae | NA | 0 1 0 0 0 |
| 4915138 | Bacteria | Actinomycetota | Actinomycetes | Micrococcales | Micrococcaceae | Arthrobacter | 0 1 1 0 1 |
| 4121904 | Bacteria | Bacteroidota | Flavobacteriia | Flavobacteriales | Weeksellaceae | Empedobacter | 0 1 0 0 1 |
| 3564189 | Bacteria | Actinomycetota | Actinomycetes | Mycobacteriales | Gordoniaceae | NA | 0 1 1 0 0 |
| 3510629 | Bacteria | Actinomycetota | Actinomycetes | Mycobacteriales | Gordoniaceae | Gordonia | 0 1 1 0 0 |
| 2926351 | Bacteria | Bacteroidota | Sphingobacteriia | Sphingobacteriales | Sphingobacteriaceae | Pedobacter | 0 1 0 0 1 |
| 2697733 | Bacteria | Pseudomonadota | Gammaproteobacteria | Enterobacterales | Enterobacteriaceae | Kosakonia | 0 1 0 0 0 |
| 2218173 | Bacteria | Bacteroidota | Bacteroidia | Bacteroidales | Prevotellaceae | NA | 0 1 0 0 1 |
| 2022135 | Bacteria | Bacteroidota | Bacteroidia | Bacteroidales | Prevotellaceae | Prevotella | 0 1 0 0 1 |
| 1697606 | Bacteria | Bacillota | Bacilli | Lactobacillales | Lactobacillaceae | Lactobacillus | 0 1 0 1 0 |
| 1436449 | Bacteria | Bacillota | Tissierellia | Tissierellales | Peptoniphilaceae | NA | 0 1 0 0 0 |
| 642730 | Bacteria | Fusobacteriota | Fusobacteriia | Fusobacteriales | Leptotrichiaceae | NA | 0 1 0 0 1 |
| 631631 | Bacteria | Pseudomonadota | Gammaproteobacteria | Enterobacterales | Morganellaceae | NA | 0 1 0 1 1 |
| 565778 | Bacteria | Pseudomonadota | Gammaproteobacteria | Enterobacterales | Enterobacteriaceae | Kluyvera | 0 1 0 0 0 |
| 432979 | Bacteria | Pseudomonadota | Betaproteobacteria | Burkholderiales | | Rhizobacter | 0 1 0 0 0 |
| 396124 | Bacteria | Bacteroidota | Flavobacteriia | Flavobacteriales | Weeksellaceae | Cloacibacterium | 0 1 1 1 1 |
| 291712378 | Bacteria | Pseudomonadota | Gammaproteobacteria | Pseudomonadales | NA | NA | 0 0 1 0 0 |
| 291506545 | Bacteria | Pseudomonadota | Gammaproteobacteria | Pseudomonadales | Pseudomonadaceae | NA | 0 0 1 0 0 |
| 47278720 | Bacteria | Actinomycetota | Actinomycetes | Mycobacteriales | NA | NA | 0 0 1 0 0 |

| | | | | | | | |
|---|---|---|---|---|---|---|---|
| 19715001 | Bacteria | Pseudomonadota | Betaproteobacteria | Burkholderiales | Oxalobacteraceae | Massilia | 0 0 1 1 0 |
| 16075272 | Bacteria | Bacteroidota | Flavobacteriia | NA | NA | NA | 0 0 1 1 0 |
| 6114626 | Bacteria | Actinomycetota | Actinomycetes | Actinomycetales | Actinomycetaceae | Actinomyces | 0 0 1 0 0 |
| 2918686 | Bacteria | Pseudomonadota | Gammaproteobacteria | Xanthomonadales | Xanthomonadaceae | Xanthomonas | 0 0 1 0 0 |
| 1645995 | Bacteria | Bacillota | Bacilli | Lactobacillales | Aerococcaceae | NA | 0 0 1 0 0 |
| 1443713 | Bacteria | Actinomycetota | Actinomycetes | Propionibacteriales | Propionibacteriaceae | Tessaracoccus | 0 0 1 1 0 |
| 1070726 | Bacteria | Actinomycetota | Actinomycetes | Micrococcales | Microbacteriaceae | Curtobacterium | 0 0 1 0 0 |
| 911676 | Bacteria | Actinomycetota | Actinomycetes | Actinomycetales | Actinomycetaceae | Schaalia | 0 0 1 0 0 |
| 195987 | Bacteria | Deinococcota | Deinococci | Thermales | NA | NA | 0 0 1 0 0 |
| 195987 | Bacteria | Deinococcota | Deinococci | Thermales | Thermaceae | NA | 0 0 1 0 0 |
| 155510 | Bacteria | Pseudomonadota | Gammaproteobacteria | Xanthomonadales | Rhodanobacteraceae | Dyella | 0 0 1 0 1 |
| 16075272 | Bacteria | Bacteroidota | Flavobacteriia | Flavobacteriales | NA | NA | 0 0 0 1 0 |
| 4908830 | Bacteria | Pseudomonadota | Alphaproteobacteria | Sphingomonadales | Sphingomonadaceae | Sphingobium | 0 0 0 1 0 |
| 2813982 | Bacteria | Actinomycetota | Actinomycetes | Micrococcales | Dermabacteraceae | NA | 0 0 0 1 0 |
| 545867 | Bacteria | Actinomycetota | Actinomycetes | Nakamurellales | Nakamurellaceae | Nakamurella | 0 0 0 1 0 |
| 361682 | Bacteria | Actinomycetota | Actinomycetes | Micrococcales | Beutenbergiaceae | NA | 0 0 0 1 0 |
| 268222 | Bacteria | Bacteroidota | Bacteroidia | Bacteroidales | Tannerellaceae | NA | 0 0 0 1 0 |
| 7768137 | Bacteria | Actinomycetota | Actinomycetes | Geodermatophilales | NA | NA | 0 0 0 0 1 |
| 7768137 | Bacteria | Actinomycetota | Actinomycetes | Geodermatophilales | Geodermatophilaceae | NA | 0 0 0 0 1 |
| 2926982 | Bacteria | Actinomycetota | Actinomycetes | Micrococcales | Dermacoccaceae | NA | 0 0 0 0 1 |
| 2150050 | Bacteria | Actinomycetota | Actinomycetes | Geodermatophilales | Geodermatophilaceae | Blastococcus | 0 0 0 0 1 |
| 2096533 | Bacteria | Bacillota | Bacilli | Lactobacillales | Streptococcaceae | Lactococcus | 0 0 0 0 1 |
| 1167029 | Bacteria | Actinomycetota | Actinomycetes | Micrococcales | Ornithinimicrobiaceae | Serinicoccus | 0 0 0 0 1 |
| 873653 | Bacteria | Bacillota | Bacilli | Lactobacillales | Carnobacteriaceae | NA | 0 0 0 0 1 |
| 301495 | Bacteria | Pseudomonadota | Betaproteobacteria | Burkholderiales | Sphaerotilaceae | Caldimonas | 0 0 0 0 1 |
| 185754 | Bacteria | Bacteroidota | Cytophagia | Cytophagales | Hymenobacteraceae | Pontibacter | 0 0 0 0 1 |
| 166985 | Bacteria | Bacillota | Clostridia | Eubacteriales | Eubacteriales Family XIII. Incertae Sedis | NA | 0 0 0 0 1 |

### 1.3 Classification models with functional profiles

#### 1.3.1 Grid Search Parameters

| **Parameter** | **Values** |
|---|---|
| Decision Tree and Extra Trees Classifier | |
| Criterion | entropy, gini index |
| Number of estimators | 10, 50, 100, 300, 500, 750, 1200 |
| Max depth | 3, 5, 8, 10, 12, 15, 20, 30, 35, 40 |
| Support Vector Classifier | |
| Kernel | linear, poly, rbf |
| Degree | 2, 3, 4, 5, 6, 7, 8 |
| K-nearest neighbors Classifier | |
| Number of neighbors | 2, 3, 4, 5, 6, 7, 8, 9, 10, 11, 12, 13, 14, 15 |
| Weights | uniform, distance |
| Multi-Layer Perceptron Classifier | |
| Hidden layer sizes | (100,), (200,), (50, 50), (20, 20, 20) |
| Activation function | relu, tanh |
| Batch size | auto, 50, 100 |
| Solver | adam, sgd |
| Maximum iterations | 1000, 3000, 5000 |
| Early stopping | False, True |
| Learning rate | constant, adaptive |

**Table S3.** Parameter values for different classifiers

#### 1.3.2 Grid Search Top Results

| Model | Accuracy | Balanced Accuracy | F1 Score |
|---|---|---|---|
| PROKKA_MLP | 0.764 | 0.726 | 0.73 |
| PROKKA_VC(soft) | 0.727 | 0.654 | 0.651 |
| Mifaser4_SVC | 0.745 | 0.663 | 0.63 |
| Mifaser4_MLP | 0.745 | 0.642 | 0.621 |
| UNIPROT_VC(hard) | 0.691 | 0.643 | 0.617 |
| UNIPROT_RandomForest | 0.691 | 0.636 | 0.601 |
| KEGG_MLP | 0.691 | 0.627 | 0.595 |
| PROKKA_VC(hard) | 0.655 | 0.605 | 0.588 |
| UNIPROT_MLP | 0.691 | 0.606 | 0.58 |
| Mifaser3_VC(soft) | 0.691 | 0.614 | 0.578 |
| Mifaser3_SVC | 0.655 | 0.606 | 0.578 |
| UNIPROT_ExtraTrees | 0.655 | 0.596 | 0.57 |
| Mifaser3_VC(hard) | 0.655 | 0.612 | 0.563 |
| KEGG_ExtraTrees | 0.691 | 0.592 | 0.562 |
| KEGG_VC(soft) | 0.618 | 0.556 | 0.559 |

**Table S4.** Top 15 performing models based on F1 Score without 5-fold cross-validation. Refer to the full document to see all results.

##### 1.3.3 Performance by city

We evaluated the performance of the best models for each annotation tool stratified by city. For the evaluation, we used 5-fold cross-validation and evaluated using F1 macro score.

| City | Mifaser4 | KEGG | Prokka |
|---|---|---|---|
| AKL | 0.774 | 0.767 | 0.774 |
| BER | 0.529 | 0.489 | 0.539 |
| BOG | 0.928 | 0.811 | 0.805 |
| DEN | 0.742 | 0.823 | 0.865 |
| DOH | 0.593 | 0.509 | 0.594 |
| ILR | 0.753 | 0.773 | 0.776 |
| LIS | 0.708 | 0.667 | 0.701 |
| NYC | 0.802 | 0.749 | 0.684 |
| SAC | 0.631 | 0.682 | 0.623 |
| TOK | 0.733 | 0.768 | 0.749 |
| BAL | 0.491 | 0.491 | 0.571 |
| MIN | 0.496 | 0.496 | 0.496 |
| SAN | 0.488 | 0.539 | 0.641 |
| SAO | 0.811 | 0.701 | 0.726 |
| VIE | 0.488 | 0.489 | 0.489 |
| ZRH | 0.489 | 0.485 | 0.486 |
| Average | 0.654 | 0.64 | 0.657 |

**Table S5.** Performance by city using F1 score

### 2 MOST ABUNDANT GENERA AND PHYLA

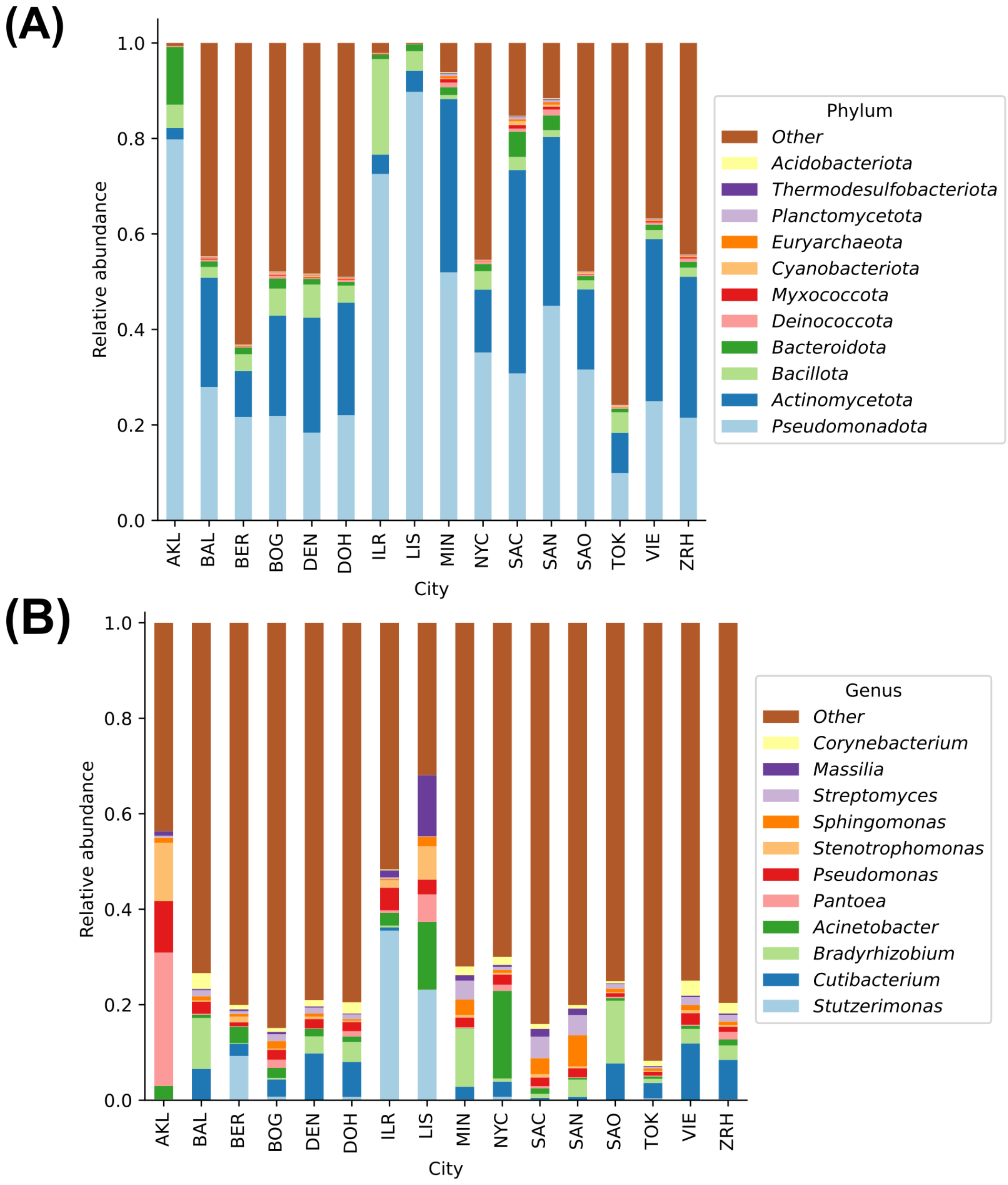


Figure S1: Relative read abundance of the most abundant microbial taxa per city at the phylum and species level, using NCBI Taxonomy as reference. **(A)** The most dominant phyla are within the Bacteria domain, with the exception of Euryarchaeota, which are archaeal organisms. The four most abundant phyla identified are consistent with the results of the urban metagenomic samples from Danko et al. (2021). **(A)** The most representative genera belong to the phylum Pseudomonadota, with the exception of Cutibacterium, Streptomyces and Corynebacterium, which belong to Actinomycetota. More than 50% of the genera found in Auckland (AKL) and Lisbon (LIS) belong to one of the eleven genera represented in the plot, reflecting the low alpha diversity found in these cities as shown in Figure 2a in the main article.

### 3 MULTI-VARIABLE LOGISTIC REGRESSION MODEL

We aim to set up a classification model starting from a set of data consisting in 365 items, where for each item we have 20,448 attributes. Each item should be classified in one of the following 16 classes: AKL, BAL, BER, BOG, DEN, DOH, ILR, LIS, MIN, NYC, SAC, SAN, SAO, TOK, VIE, ZRH.

The classification model consists in selecting $J$ of the 20,448 attributes, performing a principal component analysis on the natural logarithm of the $J$ selected attributes, and taking the first $N$ principal components as input for the multi-variable logistic regression classification algorithm.

We applied this classification scheme using several values for $J$ and $N$ and observed that classification accuracy can reach up to 88 percent when using, for example $J = 5,000$ attributes and $N = 70$ principal components. Additionally, we found that the accuracy achieved by this classification model does not improve by taking larger values for $J$.

A more detailed description of the classification scheme under consideration is provided below.

1. Take the first $J$ attributes from the total of 20,448 attributes of each of the 365 items. More precisely, consider the matrix $X$ of dimensions $365 \times J$. Each row of $X$ represents one of the 365 items to be classified with the help of the first $J$ attributes for each item. However, the attributes considered in $X$ are the natural logarithm of the original data. This is to say that the elements of $X$ are of the form $\log(d_{ij} + 1)$ were $d_{ij}$ is an entry in the original data set. It is important to note that we did not standardize the entries in matrix $X$.
2. Compute the matrix $C$ which is the correlation matrix of $X$. We call $U$ the matrix whose columns are the eigenvalues of $C$ ordered according to the decreasing order of the corresponding eigenvalues. Each column of $U$ correspond to a principal component of the transformed data matrix $X$. The quotient of the sum of the first $N$ eigenvalues and the sum of the total $J$ eigenvalues is the percentage of the variation of the data explained by the first $N$ principal components. For a given value of $J$ we choose $N$ in such a way that with the first $N$ principal components we obtain the maximum accuracy (see figure S5). With this $N$, we explain about 91% of the variation of the data.
3. The next step is to compute the matrix $Z = X \times U$ which gives the coordinates relative to the principal components of the 365 items from which we are setting up the classification model (see figure S2).
4. Each row of $Z$ has a label serving as an identifier to the class of which the corresponding item belongs to.
5. From matrix $Z$ we take a random sample $Z_{\text{tr}}$ of size 274 to be fed to a calcification algorithm. The complement of $Z_{\text{tr}}$ relative to $Z$ we call $Z_{\text{tst}}$ and will serve to test the accuracy of the classification algorithm. As classification algorithm we used the multi-variable logistic regression algorithm.
6. Once a classification model has been produced in step 5, we tested its accuracy by computing the percentage of correct classifications for each of the $91 = 365 - 274$ items in matrix $Z_{\text{tst}}$ (see figure S4).
7. Steps 5 and 6 were repeated 500 times. Each time these two steps were performed we obtained an index of the accuracy classification model. The statistical behavior of these 500 indices inform us as to what can we expect of the classification model here considered when applying it in practice.

We let $J$ take the values 2,500, 5,000, 7,500, 10,000 and 12,500. For each of these values for $J$ we performed steps 5 and 6 a total number of 500 times. The distribution of the performance index are reported in table S6 and in figure S3. We notice that the best results are obtained when $J = 10,000$ since in this

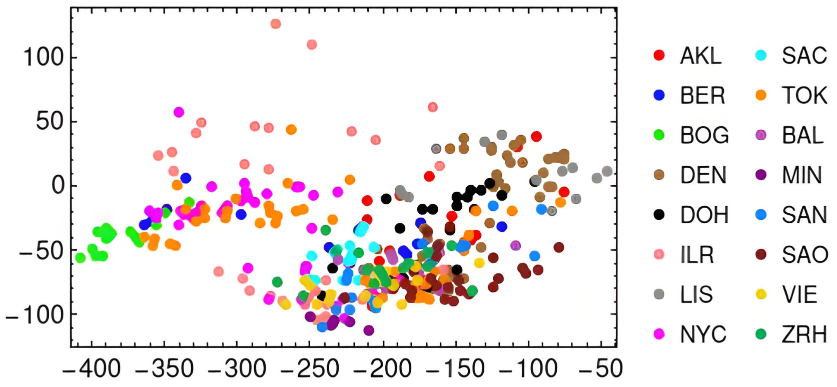


Figure S2: Here we see the distribution of the data when plotted against the first two principal components when taken the first $J = 5,000$ data attributes.

case we have a mean performance of 90% of correct classifications of the elements in the random sample of size 91 taken from the population of 365 items. A comparison of the performance of this classification model for the above five values of $J$ suggest that the performance does not improve as $J$ takes larger values. Moreover, we did not go beyond the value of 12,500 for $J$ due to the limited computation capability of a typical personal computer. For each value of $J$ we ran the classification model for distinct values of $N$ and selected the value of $N$ with the highest accuracy. Finally, this classification scheme was put to work in the Mathematica software system and we found that while the multi-variable logistic regression algorithm performed satisfactorily, other algorithms for classification did not give as good results.

| $J$ | $N$ | % | accuracy |
|---|---|---|---|
| 2,500 | 80 | 92 | 0.88 |
| 5,000 | 70 | 91 | 0.88 |
| 7,500 | 80 | 92 | 0.88 |
| 10,000 | 110 | 91 | 0.90 |
| 12,500 | 50 | 79 | 0.88 |

**Table S6.** In this table we report the accuracy achieved by the classification model here considered for distinct values of $J$ an $N$. In the third column we report the percentage of the variation of the data explained by the first $N$ principal components.

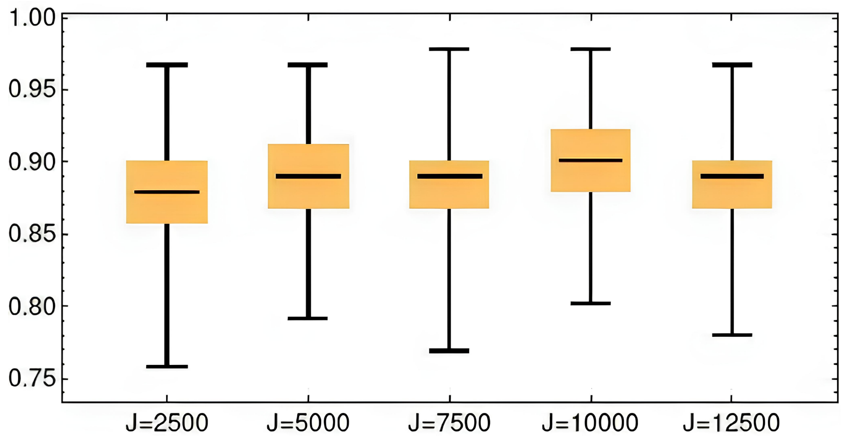


Figure S3: Box and whisker plot of 500 runs for distinct values of $J$ and for the best value of $N$.

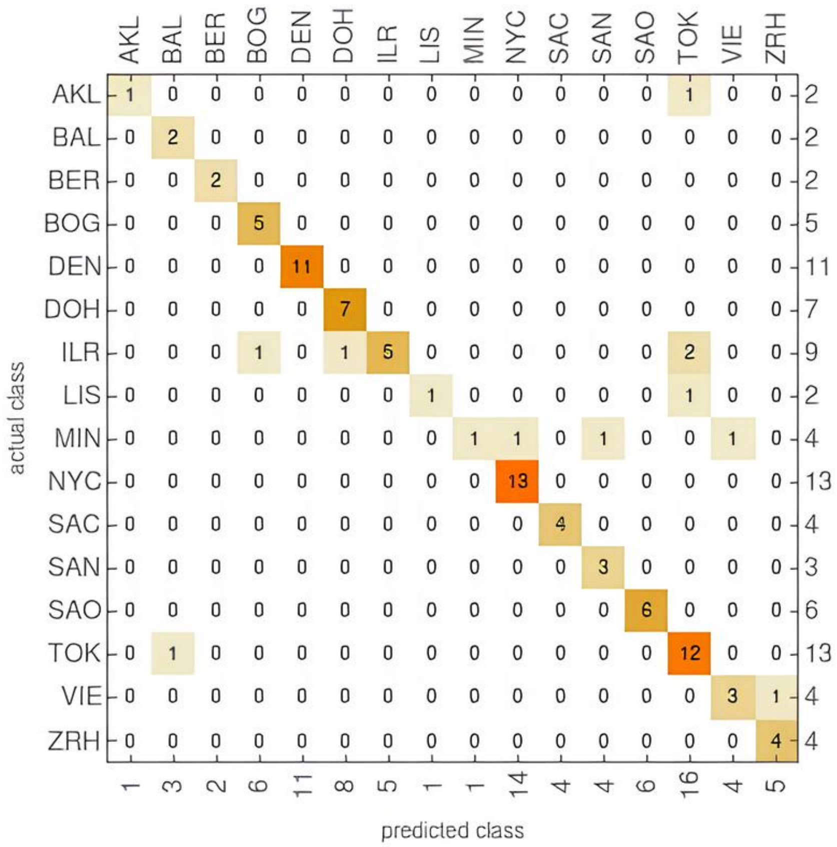


Figure S4: Confusion matrix for an accuracy of 0.879, when $J = 5,000$.

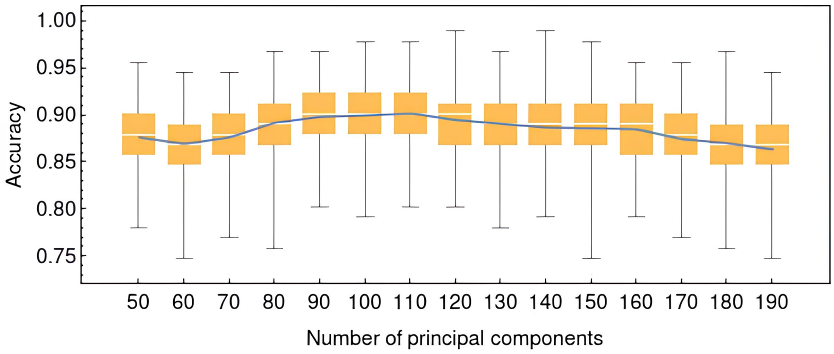


Figure S5: For $J = 10,000$ we have the performance of the classification model for distinct values of $N$. A maximum accuracy occurs when $N = 110$.
